# Complete suspension culture of human induced pluripotent stem cells supplemented with suppressors of spontaneous differentiation

**DOI:** 10.1101/2023.10.08.561419

**Authors:** Mami Matsuo-Takasaki, Sho Kambayashi, Yasuko Hemmi, Tamami Wakabayashi, Tomoya Shimizu, Yuri An, Hidenori Ito, Kazuhiro Takeuchi, Masato Ibuki, Terasu Kawashima, Rio Masayasu, Manami Suzuki, Naoki Nishishita, Yoshikazu Kawai, Masafumi Umekage, Tomoaki M Kato, Michiya Noguchi, Koji Nakade, Yukio Nakamura, Tomoyuki Nakaishi, Masayoshi Tsukahara, Yohei Hayashi

## Abstract

Human induced pluripotent stem cells (hiPSCs) are promising resources for producing various types of tissues in regenerative medicine; however, the improvement in a scalable culture system that can precisely control the cellular status of hiPSCs is needed. Utilizing suspension culture without microcarriers or special materials allows for massive production, automation, cost-effectiveness, and safety assurance in industrialized regenerative medicine. Here, we found that hiPSCs cultured in suspension conditions with continuous agitation without any microcarriers or extracellular matrix components were more prone to spontaneous differentiation than those cultured in conventional adherent conditions. Adding PKCβ and Wnt signaling pathway inhibitors in the suspension conditions suppressed the spontaneous differentiation of hiPSCs into ectoderm and mesendoderm, respectively. In these conditions, we successfully completed the culture processes of hiPSCs including the generation of hiPSCs from peripheral blood mononuclear cells with the expansion of bulk population and single-cell sorted clones, long-term culture with robust self-renewal characteristics, single-cell cloning, direct cryopreservation from suspension culture and their successful recovery, and efficient mass production of a clinical-grade hiPSC line. Our results demonstrate that precise control of the cellular status in suspension culture conditions paves the way for their stable and automated clinical application.

## Introduction

Human induced pluripotent stem cells (hiPSCs) are promising resources for various types of tissues in regenerative medicine (Takahashi *et al*, 2007; Yu *et al*, 2007). To enable cell therapy from hiPSCs, the development of a large-scale manufacturing system is essential because massive cell numbers are required to compose transplantable cells which are enough to rescue the desired physiological function (Chen *et al*, 2014; Kim & Kino-Oka, 2020; Tannenbaum & Reubinoff, 2022). In general, hiPSCs are believed to possess their scaffold dependency and are cultured under adhesion and monolayer culture conditions (Hayashi & Furue, 2016; Xu *et al*, 2001). However, utilizing suspension culture without microcarriers or special materials allows for massive production, automation, cost-effectiveness, and safety assurance in industrialized regenerative medicine.

Several attempts have been made to develop suspension culture technologies enabling rapid and large-scale preparation of hiPSCs (Amit *et al*, 2010; Amit *et al*, 2011; Dang *et al*, 2004; Elanzew *et al*, 2015; Horiguchi & Sakai, 2016; Hunt *et al*, 2014; Ibuki *et al*, 2019; Kehoe *et al*, 2010; Kim *et al*, 2019; Krawetz *et al*, 2010; Kwok *et al*, 2018; Lam *et al*, 2016; Lipsitz *et al*, 2018; Nath *et al*, 2017; Oh *et al*, 2009; Olmer *et al*, 2012; Rohani *et al*, 2020; Shafa *et al*, 2012; Singh *et al*, 2010; Steiner *et al*, 2010; Wang *et al*, 2013; Zweigerdt *et al*, 2011) (Summarized in Table 1). These studies have achieved long-term culture and/or mass expansion of hiPSCs and/or human embryonic stem cells (hESCs) in suspension conditions. However, completed processes from clonal hiPSC generation to mass production of hiPSCs based on the precise control of cell status have not yet been achieved.

**Table 1.** The list of published study on scalable suspension culture without microcarriers for hPSCs.

| Article | Medium | Additives | Cell lines tested | Scalability | Transcriptome | Single cell cloning | Direct freeze and thaw | iPSC generation |
| --- | --- | --- | --- | --- | --- | --- | --- | --- |
| Amit et al., 2010, 2011 | DMEM/F12 + KSR | Y-27632, LIF, bFGF | hESC (I3, Ie, I5, H9.2, H9, H7, H14)<br>hiPSC (iF4, J1.2.3, C3, C2, KTN7, KTN13) | $\sim 10^7$ | No | No | No | No |
| Krewetz et al., 2010 | mTeSR1 | Y-27632, Rapamycin | hESC (H9) | $\sim 10^8$ | No | No | No | No |
| Singh et al., 2010<br>Zweigerdt et al., 2011<br>Olmer et al., 2012 | KnockOut DMEM + KSR, mTeSR1 | Y-27632, bFGF | hESC (hES2, hES3, and ESI049)<br>hiPSC (hCBiPS2, hiPSOCT4eGFP) | $\sim 10^8$ | No | No | No | No |
| Steiner et al., 2010 | Neurobasal medium + KSR | Y-27632, bFGF, Activin A, Fibronectin, Gelatin, BDNF, NT3, NT4, Nutridoma-CS | hESC (HES1, HES2, H7) | $\sim 10^6$ | No | No | No | No<br>(Of note, 3 hESC lines derivation) |
| Wang et al., 2013 | TeSR-E8 | Y-27632 | hiPSC (BC1, TNC1) | $\sim 10^8$ | No | No | No | No |
| Hunt et al., 2014 | mTeSR1 | Y-27632 | hESC (H1, H9) | $\sim 10^7$ | No | No | No | No |
| Elanzew et al., 2015 | mTeSR1, TeSR-E8 | Y-27632 | hESC (H9) and hiPSC (iLB-C-31f-r1) | $\sim 10^7$ | No | No | No | No |
| Horiguchi et al., 2016; Ibuki et al., 2019 | TeSR-E8, mTeSR1 | Y-27632, KSR, Albumax, LPA, SIP, | hiPSC (TkDN4-M, TkDA3-4, 201B7) | $\sim 10^6$ | No | No | No | No |
| Nath et al., 2017 | Dialyzed DMEM/F-12 | Y-27632, bFGF, TGF-b1 | hiPSC (Tic) | $\sim 10^8$ | No | No | No | No |
| Kwok et al, 2018 | mTeSR1, StemMACS | Y-27632 | hiPSC (AR1034ZIMA hiPSC clone1, FS hiPSC clone2) | $\sim 10^9$ | No | No | No | No |
| Lipsitz et al., 2018 | Nutristem, DMEM/F12 + KSR | Y-27632, LIF, TGFβ1, FGF2, CHIR99021, SP600125, BIRB796, Gö6983 | hESC (H9, HES2, WIB) and hiPSC (C1.15) | $\sim 10^7$ | No | No | No | No |
| Rohani et al., 2020 | mTeSR1, RSeT | Y-27632, Rapamycin | hESC (H1, H9) | $\sim 10^7$ | Yes | No | No | No |
| Matsuo-Takasaki et al. 2024 (This study) | AK02N, AK03N, StemScale, mTeSR1 | Y-27632, IWR-1-endo, LY333531 | hiPSC (WTC11, 201B7, 1383D6, 1231A3, HiPS-NB1RGB, Ff-I14s04) | $\sim 10^9$ | Yes | Yes | Yes | Yes |

In this study, we have investigated what hampers the stable maintenance of undifferentiated cell states in suspension conditions. HiPSCs cultured in suspension conditions with continuous agitation without any microcarriers or extracellular matrix (ECM) components were more prone to spontaneous differentiation than those cultured in conventional adherent conditions. From screening of candidate molecules to suppress the spontaneous differentiation of hiPSCs, we have identified that inhibitors of PKCβ and Wnt signaling pathways suppress their differentiation into ectoderm and mesendoderm, respectively. In these conditions, we aimed to complete the processes of handling hiPSCs including the generation of hiPSCs with the expansion of bulk population and single-cell sorted clones, long-term culture with robust self-renewal characteristics, single-cell cloning, direct cryopreservation from suspension culture and their successful recovery, and efficient mass production of a clinical-grade hiPSC line.

## Results

### Suspension cultured hiPSCs are prone to spontaneous differentiation

First, we investigated whether the quality of hiPSCs in suspension and adherent conditions are equivalent or not. A hiPSC line, WTC11, was cultured in a conventional medium, StemFit AK02N (Ajinomoto, Tokyo, Japan), with continuous agitation (90 rpm) in non-adhesive cell culture plates for 2 passages (5 days during passages) were examined (Figure 1A). In suspension culture on days 5 and 10, hiPSCs formed round cell assemblies with slightly uneven surfaces (Figure 1B). Gene expression analysis with RT-qPCR revealed that the expression of differentiation markers, such as *PAX6* (ectoderm), *SOX17* (endoderm), and *T* (mesoderm) increased in suspension-cultured hiPSCs for 10 days (Figure 1C). To monitor the spontaneous differentiation at single-cell resolution, we established knock-in reporter hiPSC lines of PAX6-tdTomato and SOX17-tdTomato to visualize and quantify the expression of PAX6 and SOX17 at the protein level, respectively (Figure 1—figure supplement 1A–H). tdTomato-positive cells were clearly observed in day 10 samples in suspension-cultured hiPSCs, whereas no fluorescent positive cells were observed in adherent culture conditions (Figure 1D). Flow cytometric analysis revealed that hiPSCs in suspension-culture conditions contained non-negligible percentages of PAX6-tdTomato-positive and SOX17-tdTomato-positive cells (Figure 1E and 1F). Western blot analysis revealed that protein expression of PAX6 and SOX17 in suspension conditions was significantly increased (Figure 2—figure supplement 1A–D). The ratio of positive cells for a cell surface marker for undifferentiated hiPSCs, TRA1-60, was significantly lower in suspension-culture conditions (Figure 2—figure supplement 1E and F). These results suggest that a portion of hiPSCs are spontaneously differentiated in suspension conditions when cultured in conventional media. To examine global changes in gene expression patterns between suspension and adherent conditions, whole-transcriptomic RNA-seq experiments with statistical tests were performed. Gene Set Enrichment Analysis (GSEA) of all the genes and Gene Ontology Enrichment Analysis (GOEA) on differentially-regulated genes revealed that, in suspension conditions, many genes involved in differentiation toward various tissues and cell-cell adhesions were significantly up-regulated. In contrast, genes involved in nucleotide metabolism, hypoxic responses, and extracellular matrix organization were down-regulated significantly (Figure 1G–J). These results suggest that hiPSCs in suspension conditions are in the process of spontaneous differentiation into various cell lineages and are characterized by specific signatures of gene expression patterns.

**Figure 1.**
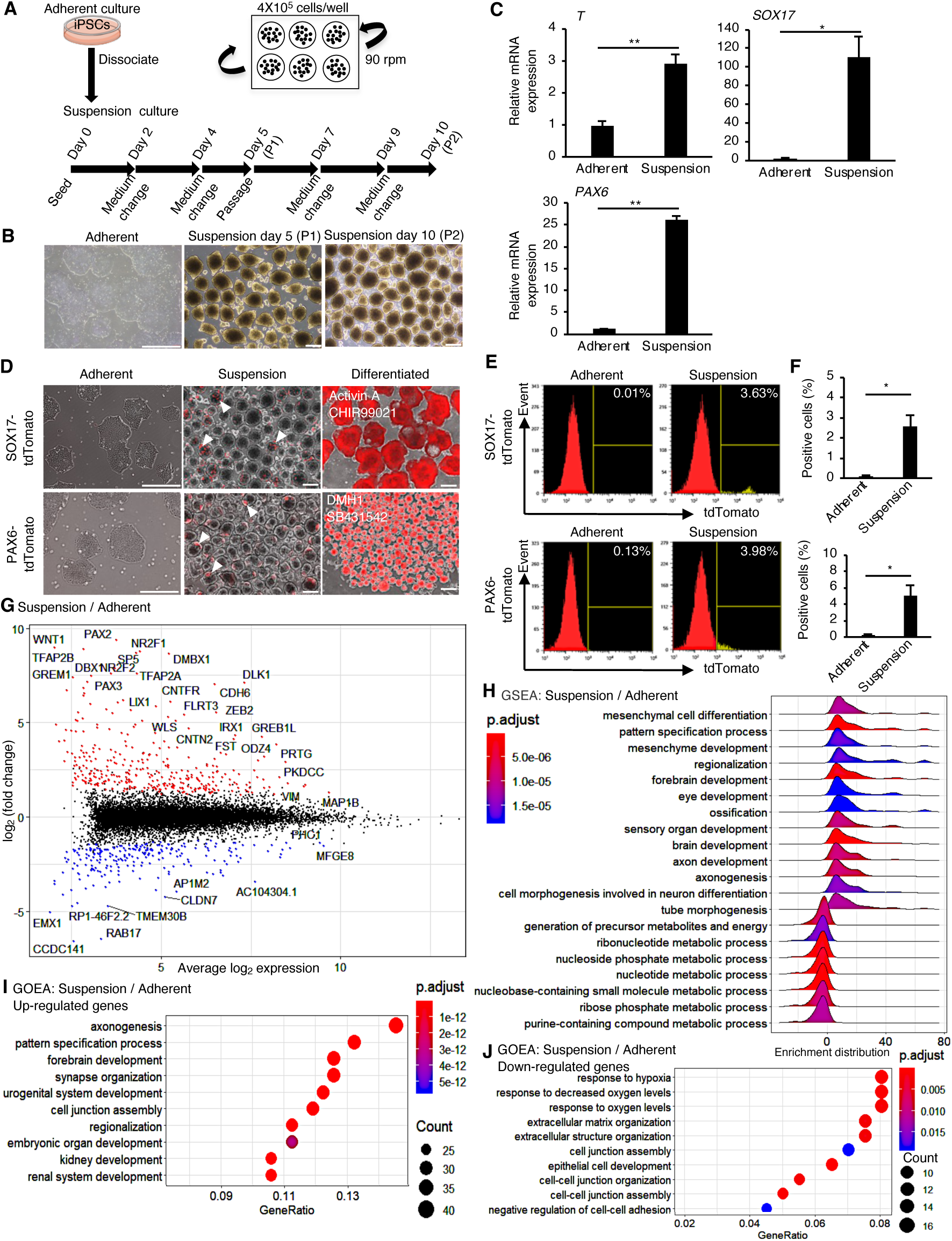
HiPSCs maintained under suspension-culture conditions undergo spontaneous differentiation. (A) Schematics representing hiPSCs in suspension-culture conditions. (B) Phasecontrast images of adherent- or suspension-cultured hiPSCs on day 5 (Passage 1 (P1)) and day 10 (Passage 2 (P2)). Scale bars: 400 µm. (C) Gene expression in hiPSCs cultured under adherent or suspension conditions on P2. Bar graphs show the mean ± SE (n=3). P-values were statistically analyzed by Student’s t-test. (D) Phase-contrast and fluorescent images of adherent or suspension-cultured reporter hiPSCs on P2. White arrowheads indicate spontaneous expression of SOX17 and PAX6 in suspension conditions. Scale bars: 400 µm. (E) Quantification of hiPSCs spontaneous differentiation with flow cytometry. (F) Averaged tdTomato-positive cell ratio (%) from flow cytometry data (mean ± SE from n=3). P-values were statistically analyzed by Student’s t-test. (G) An MA plot comparing transcriptomes between suspension- and adherent-culture from RNA-seq data. The representative gene name is shown in the plot. (H) GSEA on the gene sets of suspension-cultured hiPSCs to adherent cultures. Adjusted P-values are shown as blue to red from low to high values. (I and J) GOEA on the gene sets of suspension-cultured hiPSCs to adherent culture. Results are ranked by significance (p-adjusted value) and/or counted gene numbers.

### Wnt signaling inhibitors suppress spontaneous mesendodermal differentiation in suspension culture condition

We next investigated the factors having an inhibitory effect on hiPSCs spontaneous differentiation in suspension conditions (Figure 2A). Exogenous Wnt signaling induces the differentiation of human pluripotent stem cells into mesendoderm lineages (Nakanishi *et al*, 2009; Sumi *et al*, 2008; Tran *et al*, 2009; Vijayaragavan *et al*, 2009; Woll *et al*, 2008). Also, endogenous expression and activation of Wnt signaling in pluripotent stem cells are involved in the regulation of mesendoderm differentiation potentials (Dziedzicka *et al*, 2021). Thus, we hypothesized that adding Wnt signaling inhibitors/activators may alter the spontaneous differentiation of hiPSCs into mesendoderm. Therefore, Wnt signaling inhibitors, IWP2 or IWR-1-endo (Chen *et al*, 2009), or an activator, CHIR99021 (Ring *et al*, 2003), were added to the culture medium under suspension conditions. HiPSC aggregates treated with and without Wnt inhibitors showed similar round shapes (Figure 2B). In contrast, hiPSC aggregates treated with CHIR99021 formed heterogeneously shaped cyst-like structures, suggesting that they were largely differentiated. In samples treated with inhibitors, both T and SOX17 expression levels were significantly reduced to the level of adherent-cultured hiPSCs; however, there was only a small reduction in *PAX6* expression in the IWR-1-endo-treated condition and no reduction in the IWP2-treated condition (Figure 2C). In contrast, CHIR99021-treated cell aggregates showed markedly decreased *OCT4* and increased *T* and *SOX17* expression. Additionally, SOX17 protein expression was suppressed in hiPSCs treated with IWR-1-endo in suspension-culture conditions, although its expression increased in hiPSCs in suspension conditions with conventional culture medium compared to adherent conditions (Figure 2—figure supplement 1A and B). These results indicate that Wnt signaling inhibitors effectively suppress mesendodermal differentiation in suspension-culture, but are insufficient to suppress ectodermal differentiation.

**Figure 2.**
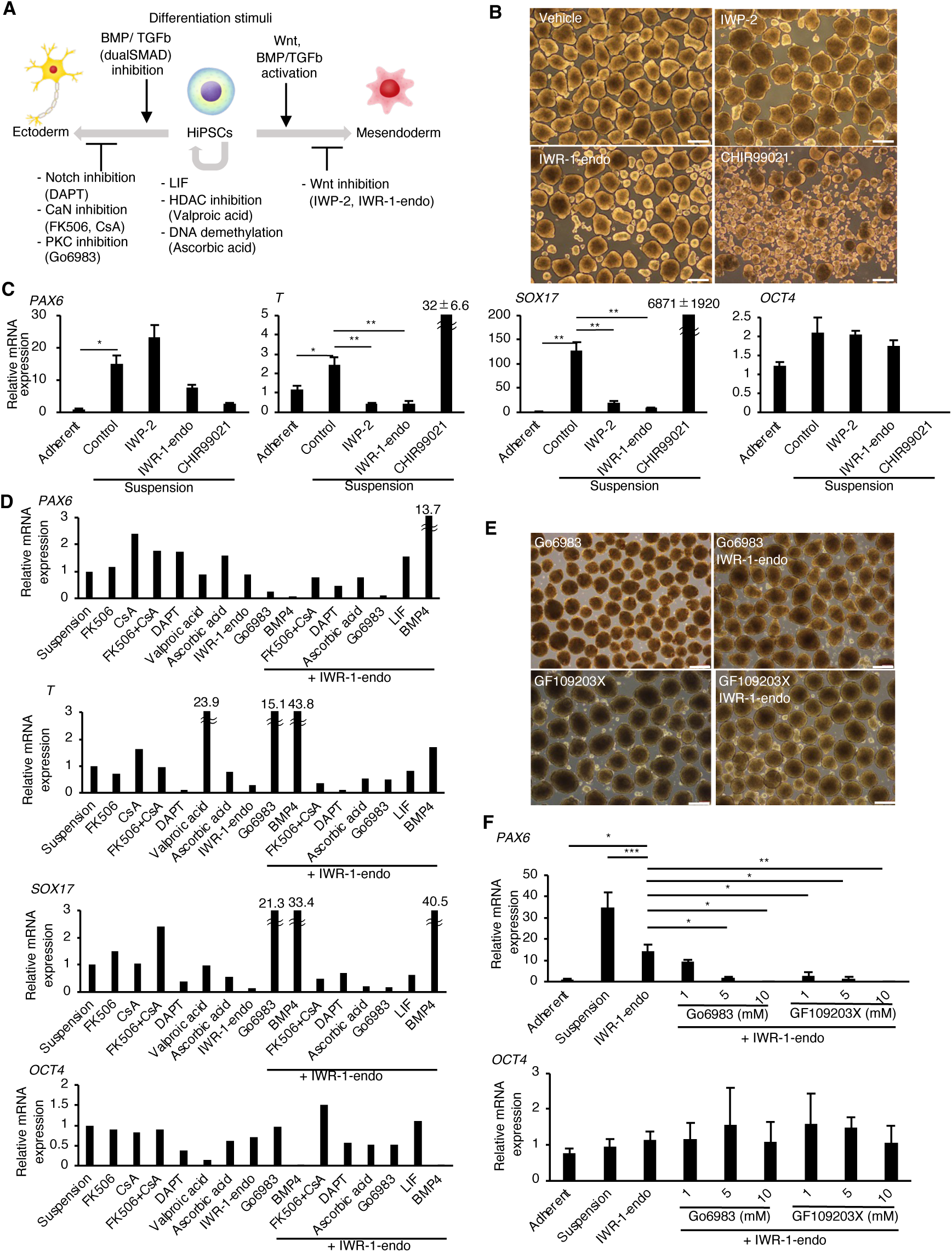
PKC inhibitors suppress spontaneous differentiation of hiPSCs into neural ectoderm in suspension-culture conditions. (A) Schematics. (B) Phase-contrast pictures of suspension-cultured hiPSCs in the presence of Wnt signaling inhibitors (IWP-2 and IWR-1-endo) or activator (CHIR99021). Scale bars: 400 µm. (C) Gene expression of hiPSCs in suspensionculture conditions with or without IWP2, IWR-1-endo or CHIR99021. RT-qPCR was performed on day 10 samples (P2). Gene expressions were normalized to GAPDH and displayed as relative fold increase to adherent-cultured samples. Bar graphs show the mean ± SE. P-values were statistically analyzed using Dunnett’s multiple comparisons test. (D) Screening of inhibitory activity of candidate molecules on neuroectoderm differentiation in suspension-cultured hiPSCs. Candidate molecules were added in combination as shown. Results are displayed as relative fold increase to suspension-cultured samples without pharmacological treatment. n=1. (E) Phase-contrast images of suspension-cultured hiPSCs on P2 in the presence of PKC inhibitors (Gö6983 or GF109203X) alone, or in combination with IWR-1-endo. Scale bars: 400 µm. (F) Suspension-cultured hiPSCs gene expression in the presence of IWR-1-endo or with combined IWR-1-endo and different doses of PKC inhibitors. Results are displayed as relative fold increase to adherent-culture. Data are presented as mean ± SE (n=3).

### PKC signal inhibitors suppress spontaneous neuroectododermal differentiation in suspension conditions

To identify molecules with inhibitory activity on neuroectodermal differentiation, hiPSCs were treated with candidate molecules in suspension conditions. We selected these candidate molecules based on previous studies related to signaling pathways or epigenetic regulations in neuroectodermal development (reviewed in (Giacoman-Lozano *et al*, 2022; Imaizumi & Okano, 2021; Sasai *et al*, 2021; Stern, 2024)) or in pluripotency safeguards (reviewed in (Hackett & Surani, 2014; Li & Belmonte, 2017; Takahashi & Yamanaka, 2016; Yagi *et al*, 2017)) (Figure 2A; listed in Supplementary Table 1). Out of the candidate molecules tested, Gö6983, a pan-PKC inhibitor (Gschwendt *et al*, 1996), and BMP4 showed strong inhibition effect on *PAX6* expression (Figure 2D). Further, simultaneous treatment with Gö6983 and IWR-1-endo decreased *PAX6*, *T*, and *SOX17* expression, while maintaining *OCT4* expression. To confirm these screening test results, the dose-dependent effect of Gö6983 or another PKCα, β, γ inhibitor GF109203X (GFX) (Toullec *et al*, 1991) on the inhibition of *PAX6* expression was observed at constant concentrations of IWR-1-endo (Figure 2E and F). These results demonstrate that the inhibition of PKC signaling pathway effectively suppresses neuroectoderm differentiation and maintain self-renewal in hiPSCs cultured in suspension conditions.

Next, we tested various PKC inhibitors to suppress neuroectodermal differentiation from hiPSCs in suspension culture using another hiPSC line, 201B7 (Figure 2—figure supplement 2A). The suppression of *PAX6* expression in hiPSCs cultured in suspension conditions was observed with PKC inhibitors, enzastaurin (Ly317615), sotrastaurin (AEB071), Ro-32-0432, Gö6983, GF109203X, and LY333531, all of which possessed PKCβ inhibition activity (Figure 2—figure supplement 2B-D); however, sotrastaurin and Ro-32-0432 also showed growth inhibition of hiPSCs.

Further, we examined the expression pattern changes in specific isoforms of PKCs in hiPSCs cultured in adherent/suspension conditions. RNA expression of PKCα (PRKCA) and PKCβ (PRKCB) was significantly up-regulated under suspension-culture conditions compared to adhesion conditions (Figure 3—figure supplement 1A). Moreover, phosphorylated PKCβ protein expression was significantly elevated (Figure 3—figure supplement 1B and C). These results suggest that elevated expression and activation of PKCβ in suspension-cultured hiPSCs could affect the spontaneous differentiation.

### Combination of inhibitors of PKCβ and Wnt signaling pathways efficiently maintains self-renewal of hiPSCs in suspension conditions

To further explore the possibility that the inhibition of PKCβ is critical for the maintenance of self-renewal of hiPSCs in the suspension culture, we evaluated the effect of LY333531, a PKCβ specific inhibitor (Jirousek *et al*, 1996). As compared to controls, a hiPSC line, WTC11, cultured in suspension conditions treated with IWR-1-endo and LY333531 formed homogenous, round, smooth-surfaced aggregates (Figure 3A). *PAX6* expression was strongly suppressed by the addition of LY333531 (Figure 3B). Furthermore, after adding IWR-1-endo, the inhibitory effect of *PAX6* expression was further enhanced, and simultaneously, *OCT4* expression was restored to the same level as in the adherent-culture. PAX6 protein expression was also suppressed in hiPSCs treated with IWR-1-endo and LY333531 in suspension-culture conditions while its expression increased in suspension conditions with conventional culture medium compared to adherent-conditions (Figure 2—figure supplement 1C and D). The ratio of TRA-1-60-positive cells was higher in suspension-culture conditions supplemented with IWR-1-endo and LY333531 than in control conditions without these inhibitors (Figure 2—figure supplement 1E and F). These results indicate that the maintenance of suspension-cultured hiPSCs is specifically facilitated by the combination of PKCβ and Wnt signaling inhibition.

**Figure 3.**
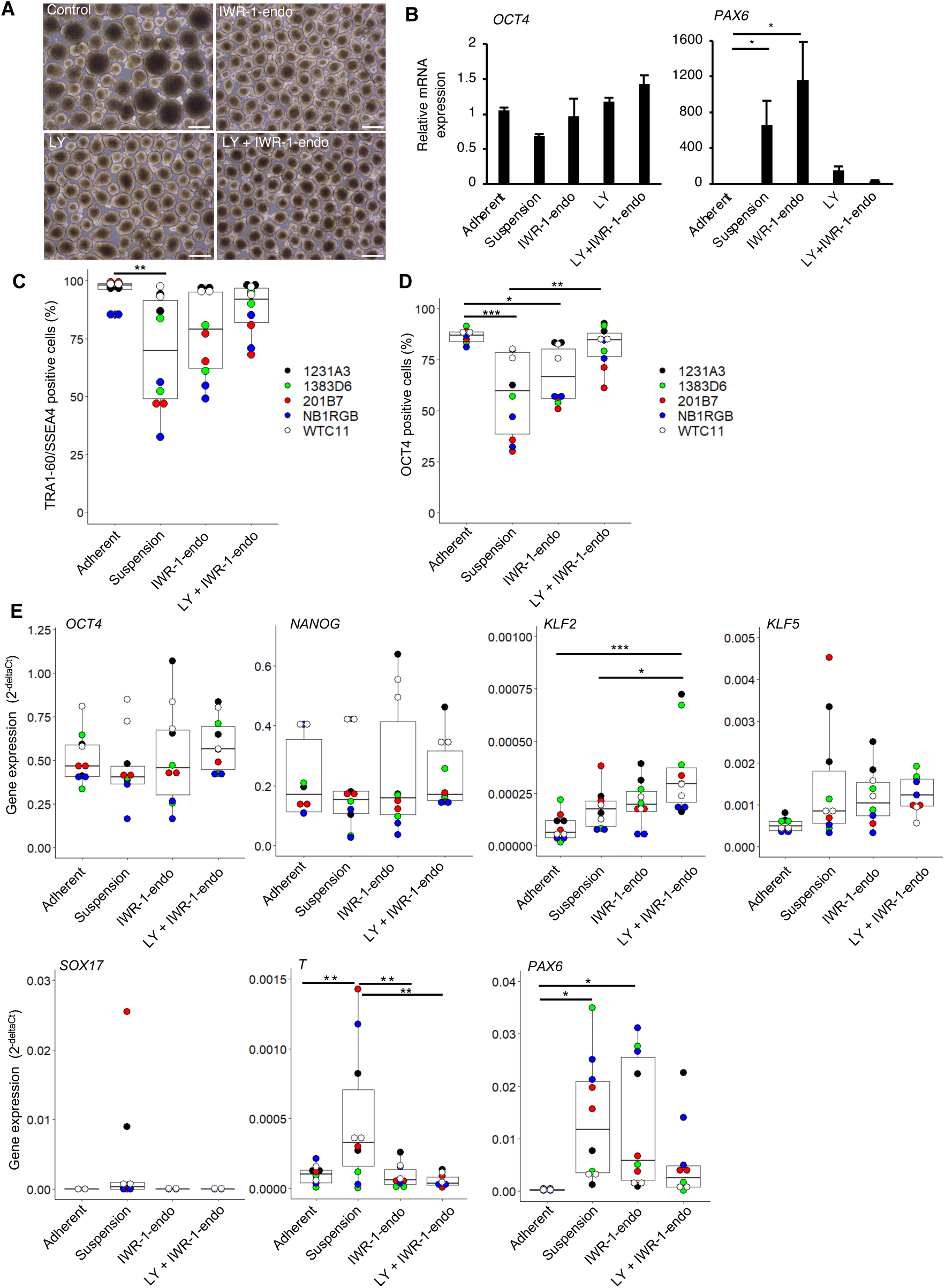
The inhibitors of WNT and PKCβ efficiently maintain the self-renewal of hiPSCs in suspension conditions. (A) Phase-contrast images of suspension-cultured hiPSCs on day 10 (2 passages) in the presence of IWR-1-endo, LY333531, or both. Scale bars: 400 µm. (B) Gene expression of suspension-cultured hiPSCs on P2 in the presence of IWR-1-endo, LY333531, or both. Data are presented as mean ± SE. P-values were statistically analyzed with Dunnett’s test. (C) After suspension culture of 5 hiPSC lines for 10 days (2 passages), WTC11, 1231A3, HiPS-NB1RGB (NB1RGB), 1383D6 and 201B7, in StemFit AK02N medium with or without IWR-1-endo and LY333531, characteristic analysis was performed. Box plots of flow cytometry for TRA-1-60 and SSEA4 double positive cells (%) are shown. Used cell lines are indicated by the different colored circles shown on the right side of the graph (n=2 x 5 cell lines). (D) Box plots of OCT4 positive cells (%) are shown (n=2 x 5 cell lines) (E) Box plots of RT-qPCR data are shown. Differentiation markers (*SOX17*, *T*, *PAX6*), undifferentiated markers (*OCT4, NANOG*), and naïve markers (*KLF2*, *KLF5*) were assessed. Statistical analysis was performed using by one-way Anova and Tukey’s test for all graphs. P values < 0.05 were considered statistically significant. *, ** or *** in the graphs indicate P<0.05, P<0.01 or P<0.001, respectively (n=2 x 5 cell lines).

To examine the reproducibility of the effect of the inhibition of PKCβ and Wnt signaling pathways on the maintenance of self-renewal of hiPSCs in the suspension culture among various hiPSCs, we evaluated the expression of self-renewal and differentiation markers among 5 different hiPSC lines, 1231A3, 1383D6, 201B7, HiPS-NB1RGB, and WTC11, simultaneously. Compared to adherent culture conditions, hiPSCs cultured in suspension conditions without chemical treatment decreased the positive ratio of TRA-1-60/SSEA4 and OCT4 and increased the expression levels of differentiation markers, *SOX17*, *T*, and *PAX6* (Figure 3C-E). These results indicated that suspension culture conditions without chemical treatment are unstable to maintain self-renewal and contain spontaneous differentiated cells. The addition of PKCβ and Wnt signal inhibitors increased the positive ratio of TRA-1-60/SSEA4 and OCT4 and decreased the expression levels of *SOX17*, *T*, and *PAX6* to the comparable level of adherent conditions. Interestingly, the expression of KLF2 and KLF5, which are also known as naïve pluripotency markers, were up-regulated in the suspension conditions treated with PKCβ and Wnt signal inhibitors.

We also examined whether the combination of PKCβ and Wnt signaling inhibition affects cell survival in suspension conditions. In this experiment, we used another PKC inhibitor, Staurosporine (Omura *et al*, 1977), which has a strong cytotoxic effect as a positive control of cell death in suspension conditions. The addition of IWR-1-endo and LY333531 for 10 days had no effects on the apoptosis while the addition of Staurosporine for 2 hours induced Annexin-V-positive apoptotic cells (Figure 3—figure supplement 2A-D). These results indicate that the combination of PKCβ and Wnt signaling inhibition has no or little effects on the cell survival in suspension conditions.

We next performed long-term culture for 10 passages in suspension conditions and compared hiPSC growth in presence of LY333531 or Gö6983. When hiPSCs were seeded at 4 × 10^5^ cells/well, the average cell number reached approximately 12-fold after 5 days under both conditions (Figure 4A and B). After 10 passages, aggregates of hiPSCs cultured in the presence of LY333531 showed a uniform spherical shape, whereas aggregates cultured in the presence of Gö6983 were heterogeneously spherical (Figure 4C). Notably, in LY333531-treated cells, OCT4-positive cell numbers were significantly higher than in Gö6983-treated samples, as determined by immunostaining (Figure 4D and E). To evaluate whether hiPSCs cultured in suspension conditions with PKCβ and Wnt signaling inhibitors for 10 passages maintain pluripotency, we performed embryoid body (EB) formation assay. These EBs contained positive cells for TUJ1, SMA, and AFP as ectodermal, mesodermal, and endodermal marker, respectively (Figure 4F). Copy number variation (CNV) array analysis showed that hiPSCs cultured long-term in the presence of PKCβ and Wnt inhibitors retained their normal human karyotype (Figure 4G). These results indicate that, for long-term culture, the inhibition of Wnt signaling and PKCβ in suspension conditions is sufficient to maintain the self-renewal, pluripotency, and genomic integrity of hiPSCs. Thus, we used the combination of IWR-1-endo and LY333531 for the rest of this study.

**Figure 4.**
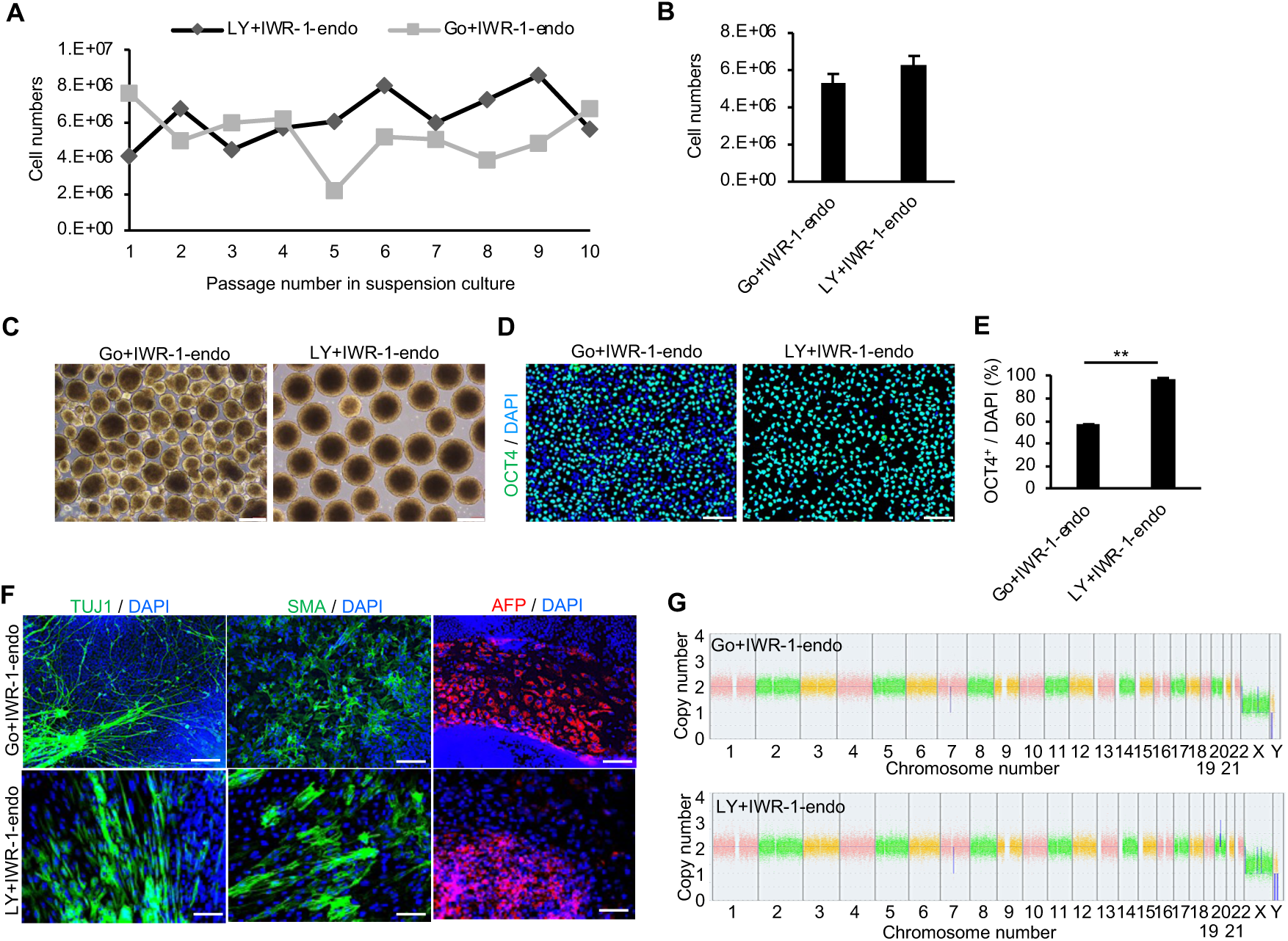
Long-term suspension culture of hiPSCs is maintained by simultaneous suppression of PKCβ and Wnt signals. (A) The number of hiPSCs at passage during long-term suspension-culture every 5 days. Cell culture was performed in the presence of IWR-1-endo plus Gö6983 or LY333531. (B) Bar graph indicating average cell numbers for 10 passages (mean ± SE). (C) Phase-contrast images of suspension-cultured hiPSCs on passage 10. Scale bars: 400 µm. (D) Immunocytochemistry of OCT4 on passage 10 samples. Scale bars: 100 µm. (E) Bar graph showing the percentages of OCT4 positive cells. Values were calculated from randomly selected three regions from immunofluorescence images. Data are presented as mean ± SE. P-value was statistically analyzed by Student’s t-test. (F) Immunocytochemistry of differentiated cells in EBs from suspension-cultured hiPSCs in the presence of IWR-1-endo; and Gö6983 or LY333531. Anti-TUJ1, -SMA and -AFP antibodies were used to detect ectoderm, mesoderm and endoderm differentiation, respectively. Scale bars:100 µm. (G) Chromosomal copy numbers detected with CNV array analysis (Karyostat assay) of suspension-cultured hiPSCs in the presence of IWR-1-endo; and Gö6983 (Upper panel) or LY333531 (Lower panel) at passage 10.

We further investigated whether the effects of PKCβ and Wnt inhibitors on suppressing hiPSCs spontaneous differentiation in suspension-culture conditions are applicable to other culture media. First, morphologies and gene expression profile of hiPSCs cultured in suspension conditions with another commercially available maintenance medium, StemScale (ThermoFisher Scientific, MA, USA), were examined (Figure 4—figure supplement 1A). A hiPSC line, WTC11, cultured in suspension treated with IWR-1-endo and LY333531 formed homogenously round and smooth-surfaced aggregates compared to controls (Figure 4—figure supplement 1B). The differentiation markers expression, which was elevated under suspension conditions, was suppressed by the simultaneous addition of PKCβ and Wnt signal inhibitors, as observed with StemFit medium (Figure 4—figure supplement 1C and D). Second, morphologies and gene expression profile of hiPSCs, cultured in suspension conditions with another commercially available maintenance medium, mTeSR1 (Stem Cell Technologies, Vancouver, Canada), were examined in 4 different hiPSC lines (1231A3, 201B7, HiPS-NB1RGB, and WTC11) simultaneously (Figure 4—figure supplement 2A). Compared to adherent culture conditions, hiPSCs cultured in suspension conditions without chemical treatment significantly decreased the positive ratio of TRA-1-60/SSEA4 and OCT4 and increased the expression levels of differentiation markers, *SOX17*, *T*, and *PAX6* (Figure 4—figure supplement 2B-D). These results indicated that suspension culture conditions using mTeSR1 medium contain spontaneous differentiated cells. The addition of PKCβ and Wnt signal inhibitors increased the positive ratio of TRA-1-60/SSEA4 and OCT4 and decreased the expression levels of SOX17, T, and PAX6 to the comparable level of adherent conditions. Together, these results suggest that conventional suspension culture methods contain spontaneous differentiating cells and that the addition of inhibitors of PKCβ and Wnt signaling pathways to conventional culture media generally suppresses spontaneous differentiation and maintains self-renewal.

### Global gene expression signatures in hPSCs supplemented inhibitors of PKCβ and Wnt signaling pathways in suspension conditions

We next examined the effect of PKCβ and Wnt inhibitors on global gene expression of hiPSCs under suspension conditions. Bulk RNA-seq data were obtained from suspension conditions in absence of inhibitors (Sus), supplemented with IWR-1-endo (IWR), LY333531 (LY), IWR-1-endo and LY333531 (IWRLY), and adherent conditions (Ad). Hierarchical clustering obtained from these data showed that LY and IWRLY were grouped closely with Ad (Figure 5A). In contrast, Sus and IWR were both grouped as discrete populations from Ad. Additionally, hierarchical clustering in gene expression among these conditions was supported by principal component 1 (PC1) in the principal component analysis (PCA) (Figure 5B). In contrast, PC2 represented genes related to the effects of specific inhibitors under these conditions. Next, we investigated the effect of LY333531 and IWR-1-endo in suspension culture. Many genes involved in pluripotency: *KLF4* and *ID1*, and epithelial cell-cell interactions: *CDH1* (E-cadherin), were significantly up-regulated in IWRLY, while many transcription factors involved in differentiation: *PAX2*, *PAX3, PAX5, PAX8*, *SP5*, *DBX1*, and *TFAP2B*, were down-regulated in IWRLY (Figure 5C). GSEA and GOEA on down-regulated genes in IWRLY showed that the expression of developmentally associated genes, whose expression was elevated in Sus, was generally reduced in IWRLY (Figure 5D and E). GOEA on up-regulated genes revealed gene sets involved in epithelial cell types (Figure 5F). As compared to Ad, genes involved in sensory system development, cell-cell adhesion, and Wnt and PI3K signaling pathways were up-regulated in IWRLY, and genes involved in nucleotide metabolism and hypoxic responses were down-regulated under IWRLY conditions (Figure 5—figure supplement 1A-D). These results suggest that PKCβ and Wnt signaling inhibitors in suspension conditions regulate global gene expression patterns to suppress spontaneous differentiation, albeit remaining expression signatures of suspension culture, possibly due to the microenvironment within the formed aggregates and physiological differences. We also extracted and analyzed individual gene expression data of pluripotency markers from RNA-seq results. Compared to adherent condition, the expression of naïve pluripotency markers, *KLF2*, *KLF4*, *KLF5*, and *DPPA3*, were up-regulated in IWRLY conditions while *OCT4* and *NANOG* were at the similar levels (Figure 5—figure supplement 2). Combined with RT-qPCR analysis data on 5 different hiPSC lines (Figure 3E), these results suggest that IWRLY conditions may drive hiPSCs in suspension conditions to shift toward naïve pluripotent states.

**Figure 5.**
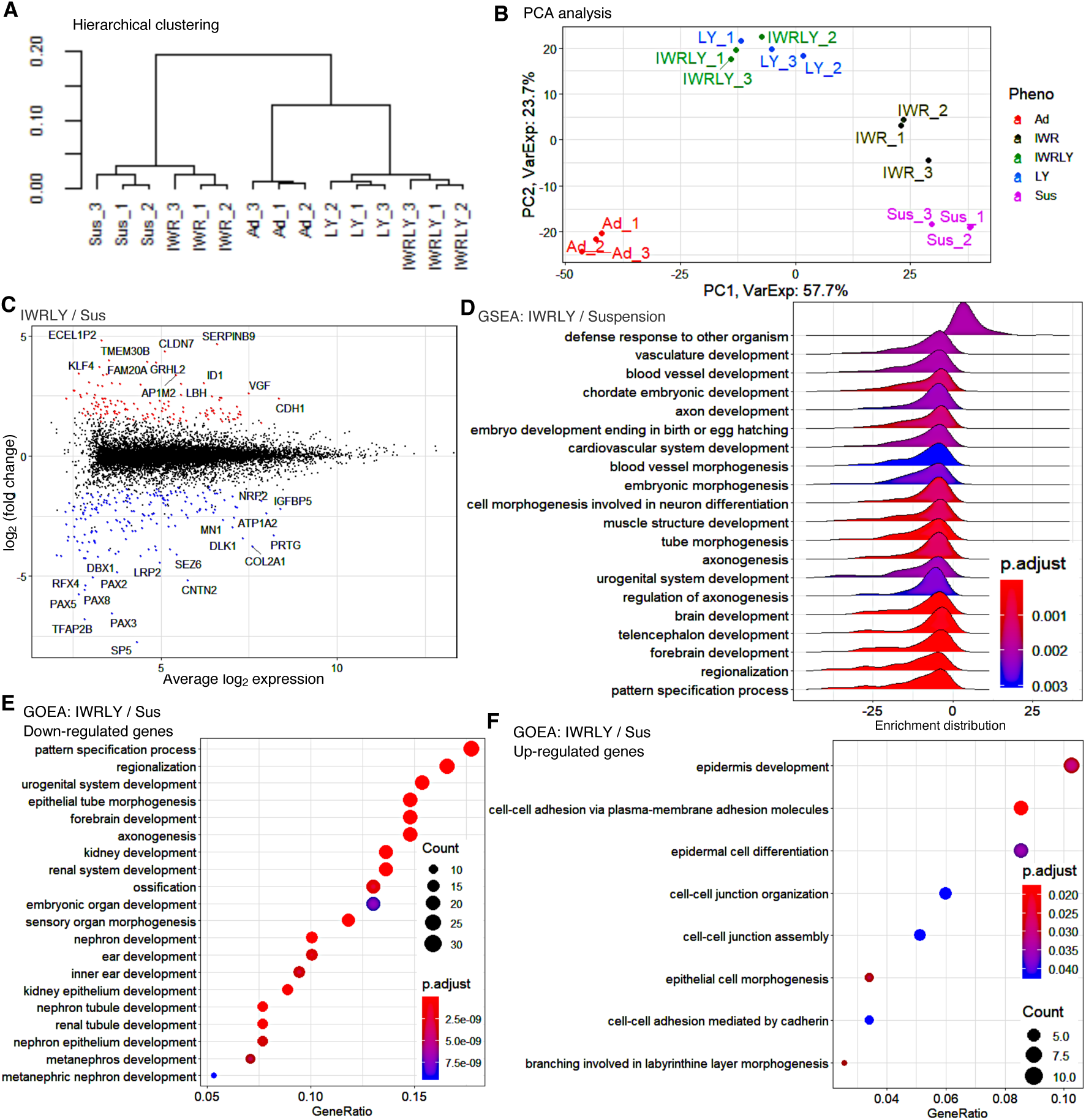
The inhibitors of WNT and PKCβ in suspension conditions efficiently suppress differentiated gene marker expression in transcriptome analysis. (A) Hierarchical clustering of adherent and suspension-cultured hiPSCs on P2 using Ward’s method from RNA-seq data (n=3 in each condition). Ad, Adherent; Sus, Suspension; IWR, IWR-1-endo; LY, LY333531; IWRLY, IWR-1-endo and LY333531. (B) Principal component analysis (PCA) plot showing clusters of samples based on similarity. Gene expression variance are displayed as PC1 and PC2. (C) MA plot comparing transcriptomes between IWRLY and Sus. (D) GSEA on the gene sets of IWRLY to Sus from these RNA-seq data. Statistically significant enrichment is shown. P-values are represented in blue to red from low to high values. (E and F) GOEA for the gene sets of IWRLY to Sus from these RNA-seq data. Analysis was performed on down-regulated genes in (E) and up-regulated genes in (F).

### Mass expansion of hiPSCs in suspension conditions supplemented with inhibitors of PKCβ and Wnt signaling pathways

For the clinical applications of hiPSCs, its homogenous mass production is required to obtain sufficient quantities. To test the feasibility of mass production under suspension-culture conditions supplemented with PKCβ and Wnt signal inhibitors, we first performed suspension-culture using a healthy-donor derived hiPSC line, 1383D6, in a 30 mL bioreactor with stirring conditions (Matsumoto *et al*, 2022) at different cell seeding densities and different stirring speeds (Figure 6—figure supplement 1A). HiPSCs steadily proliferated at 0.5 x 10^5^ - 2 x 10^5^ cells/mL of the cell seeding density, but cells hardly proliferated at 8 x 10^5^ cells/mL (Figure 6—figure supplement 1B). Since the number of total collected cells was the lowest at a seeding density of 0.5 x 10^5^ cells/mL, the seeding density of 1 x 10^5^ - 2 x 10^5^ cells/mL is considered suitable. Also, hiPSCs steadily proliferated at 50 - 150 rpm of the stirring speed, but not at 250 rpm (Figure 6— figure supplement 1C). Then, we analyzed protein expression of PAX6 and SOX17 in these cells after 3 passages with these conditions at 50 - 150 rpm of stirring speed. Suspension culture in the presence of PKCβ and Wnt inhibitors decreased the positive ratio of PAX6 and SOX17 in these reactor stirring speeds (Figure 6—figure supplement 1D). These results suggest that PKCβ and Wnt inhibitors suppressed spontaneous differentiation in bioreactor conditions at suitable stirring speeds.

Next, we examined that undifferentiated hiPSCs were efficiently maintained in these bioreactor conditions with 5 serial passages using two hiPSC lines, 1383D6 and 1231A3 (Figure 6—figure supplement 2A). These cell lines showed higher cell density in the bioreactor at each passage after 3 days (Figure 6—figure supplement 2B). Under these conditions, the concentration of glucose decreased daily (Figure 6—figure supplement 2C), whereas that of L-lactic acid increased (Figure 6—figure supplement 2D). These results indicate that the cell lines proliferated with active energy consumption. Further, RT-qPCR analysis showed no marked differences in the undifferentiated and differentiation markers expression between hiPSCs cultured in adhesion vs. suspension with PKCβ and Wnt inhibitors (Figure 6—figure supplement 2E and F). Flow cytometric analysis showed that more than 90% of these cells were positive for pluripotency marker proteins: NANOG, OCT4, and SOX2 (Figure 6—figure supplement 2G). G-band analysis of suspension-cultured hiPSCs after five passages revealed normal karyotype (Figure 6—figure supplement 2H). These results demonstrate that undifferentiated hiPSCs were efficiently maintained in the bioreactor using the culture medium supplemented with inhibitors of PKCβ and Wnt signaling pathways.

Then, we expanded hiPSCs in a large-scale culture system under perfusion conditions in the presence of IWR-1-endo and LY333531. In this experiment, clinical-grade hiPSC line, Ff-l14s04, which is derived from PBMCs of a donor carrying homozygous alleles for major HLA loci (HLA-A, HLA-B, and HLA-DR) was used (Kitano *et al*, 2022). Large-scale hiPSCs preparation using a perfusion-culture system with 320 mL bioreactor having stirred-wing (Kropp *et al*, 2016) was performed in GMP-compliant, clinical grade StemFit AK03N medium containing IWR-1-endo and LY333531 (Figure 6A). When the culture was started at 1 × 10^5^ cells/mL in 320 mL medium, hiPSCs proliferated nearly 10-fold after 3 - 4 days to produce ∼300 stock vials (1 × 10^6^ cells/vial). This large-scale culture was repeated three times (Passages 1, 2, and 3). Since the population doubling time (PDT) of this hiPSC line in adherent culture conditions is 21.8 - 32.9 hours determined at its production, this proliferation rate in this large scale suspension culture is comparable to adherent culture conditions. The frozen vials of this hiPSC lines obtained at each passage in large-scale suspension culture conditions were characterized. Cells obtained from large-scale suspension cultures formed typical hiPSC-like colonies after vials were thawed and seeded in adherent-culture conditions (Figure 6B).These samples from suspension culture conditions showed similar or higher viability (>90%) to that of adherent-culture derived vials (Figure 6C). When compared to adherent conditions, these samples from suspension culture conditions showed a similar or higher proliferation rate after thawing (Figure 6D). Flow cytometric analysis showed that over 90% of cells were positive for pluripotent cell markers expression: TRA-1-60, SSEA4, and OCT4 (Figure 6E and F). Further, G-band analysis revealed that hiPSCs retained their normal karyotype even after three passages under large-scale suspension conditions (Figure 6G). When large-scale suspension-culture derived hiPSCs were incubated in each germ layer-specific differentiation medium for 4 - 7 days, the expression of early differentiation markers for ectoderm (*PAX6* and *SOX1*), mesoderm (*T* and *PDGFRA*) and endoderm (*SOX17* and *CXCR4*) was significantly induced (Figure 6H). These cells were then directly differentiated into dopaminergic neural progenitors, cardiomyocytes, and hepatocytes to evaluate their differentiation capacity and propensity. There were no differences in the differentiation efficiency toward these lineages (Figure 6I-M). These results indicate that the characteristics and quality of clinical-grade hiPSCs cultured in large-scale suspension conditions in the presence of PKCβ and Wnt inhibitors are equivalent to those of hiPSCs maintained under adherent conditions. Taken together, we made successful in mass suspension-culture conditions of hiPSCs supplemented with Wnt and PKCβ inhibitors.

**Figure 6.**
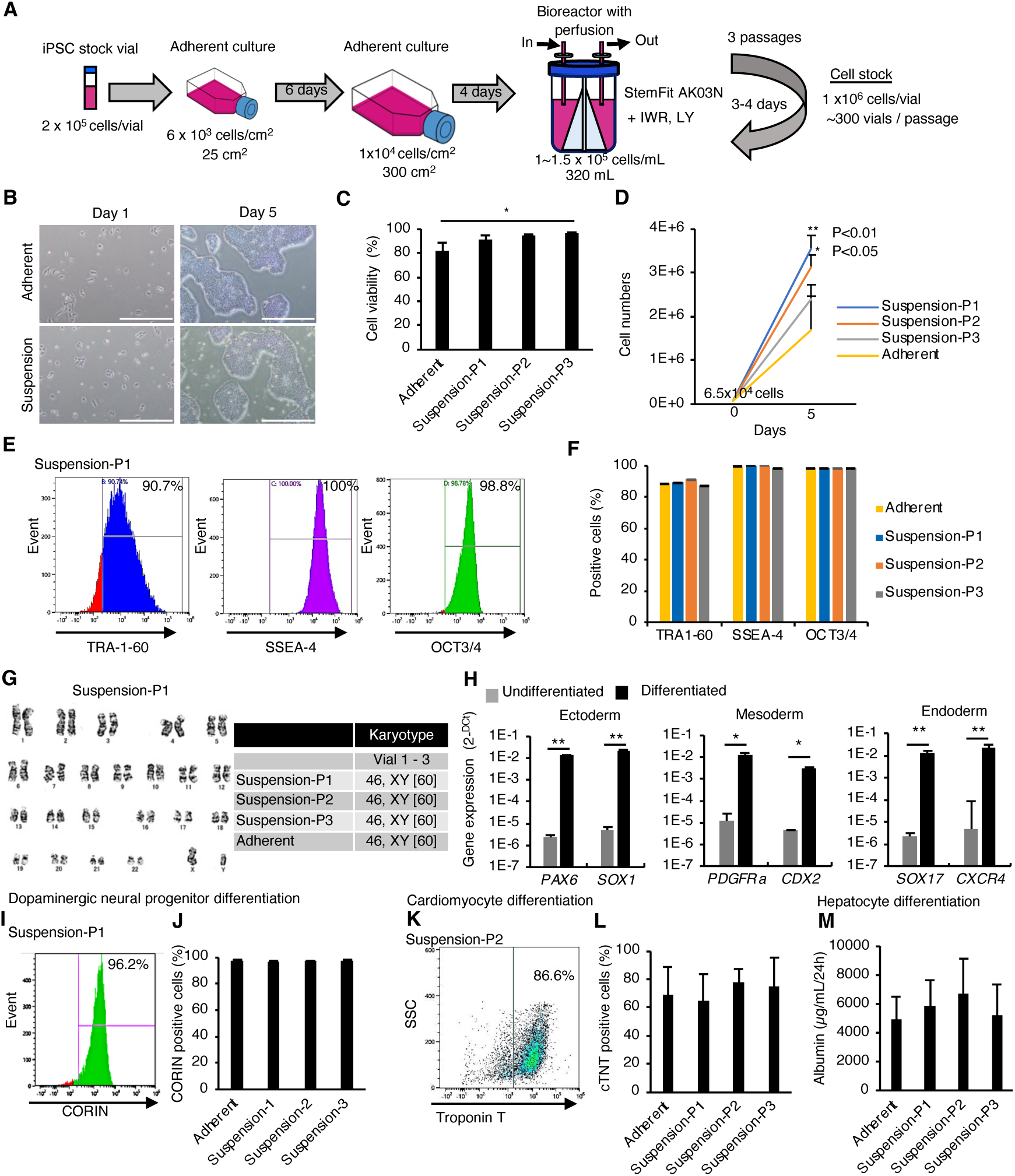
Mass suspension-culture of clinical-grade hiPSCs in the presence of PKCβ and Wnt signaling inhibitors. (A) Schematics. (B) Representative phase-contrast images of these hiPSCs. Scale bar: 1 mm. (C) Cell viability at seeding. Data are presented as mean ± SE (n=3). P-values were statistically analyzed with Dunnett’s multiple comparisons test. (D) Total cell numbers were counted on day 5 after thawing. Data were presented as mean ± SE (n=3). P-values were statistically analyzed with Dunnett’s test. (E) Representative flow cytometry data of pluripotent markers in these hiPSCs. (F) Quantification of flow cytometry data for the pluripotent markers. Data are presented as mean ± SE (n=3). (G) Karyotypes of these hiPSCs. Left: Representative pictures of G-band analysis. Right: Table of karyotype results (n=3). The numbers in brackets indicate the cell numbers examined. (H) *In vitro* differentiation from hiPSCs was assessed using RT-qPCR (mean ± SE) (n=3). P-value was statistically analyzed by Student’s t-test. (I) Representative flow cytometry data for CORIN. (J) Quantification of CORIN-positive cells (mean ± SE, n=3). (K) Representative flow cytometry data for cardiac Troponin T (cTnT). (L) Quantification of cTnT-positive cells (mean ± SE, n=3). (M) Albumin secretion levels of hepatocytes differentiated from hiPSCs (mean ± SE, n=3).

### Single-cell sorting and expansion of hiPSC subclones cultured in suspension conditions

To test the feasibility of our suspension culture method to single cell sorting, we sorted a hiPSC line, 201B7, with TRA-1-60 antibody into individual wells of a 96-well plate and attempted to establish subclones by expanding them with serial passages using StemFit AK02N medium (Figure 7A). On day 7 after single cell sorting, we counted the number of colonies to calculate the cloning efficiency (Figure 7B). The cloning efficiency in adherent culture conditions was around 30%. While the cloning efficiency in suspension culture conditions without any chemical treatment was less than 10%. The treatment of IWR-1-endo in the suspension culture conditions increased the efficiency more than 20% although the treatment of LY333531 decreased the efficiency. These results indicated that the IWR-1-endo treatment is beneficial in single-cell cloning in suspension culture conditions. On day 14, we performed passages of formed colonies and cultured in suspension conditions supplemented with IWR-1-endo LY333531. By day 28 after this single-cell sorting, we expanded single-cell-sorted hiPSC subclones in suspension culture supplemented with IWR1-endo and LY3333531. On day 28, we examined seven subclones for their cell growth and expression of OCT4 and TRA-1-60. The subclones showed round-shaped aggregates with more than three million cells and high ratios of OCT4- and TRA1-60-positive cells (Figure 7C-F). These results indicate that we have successfully derived single-cell-cloned sublines in suspension culture.

**Figure 7.**
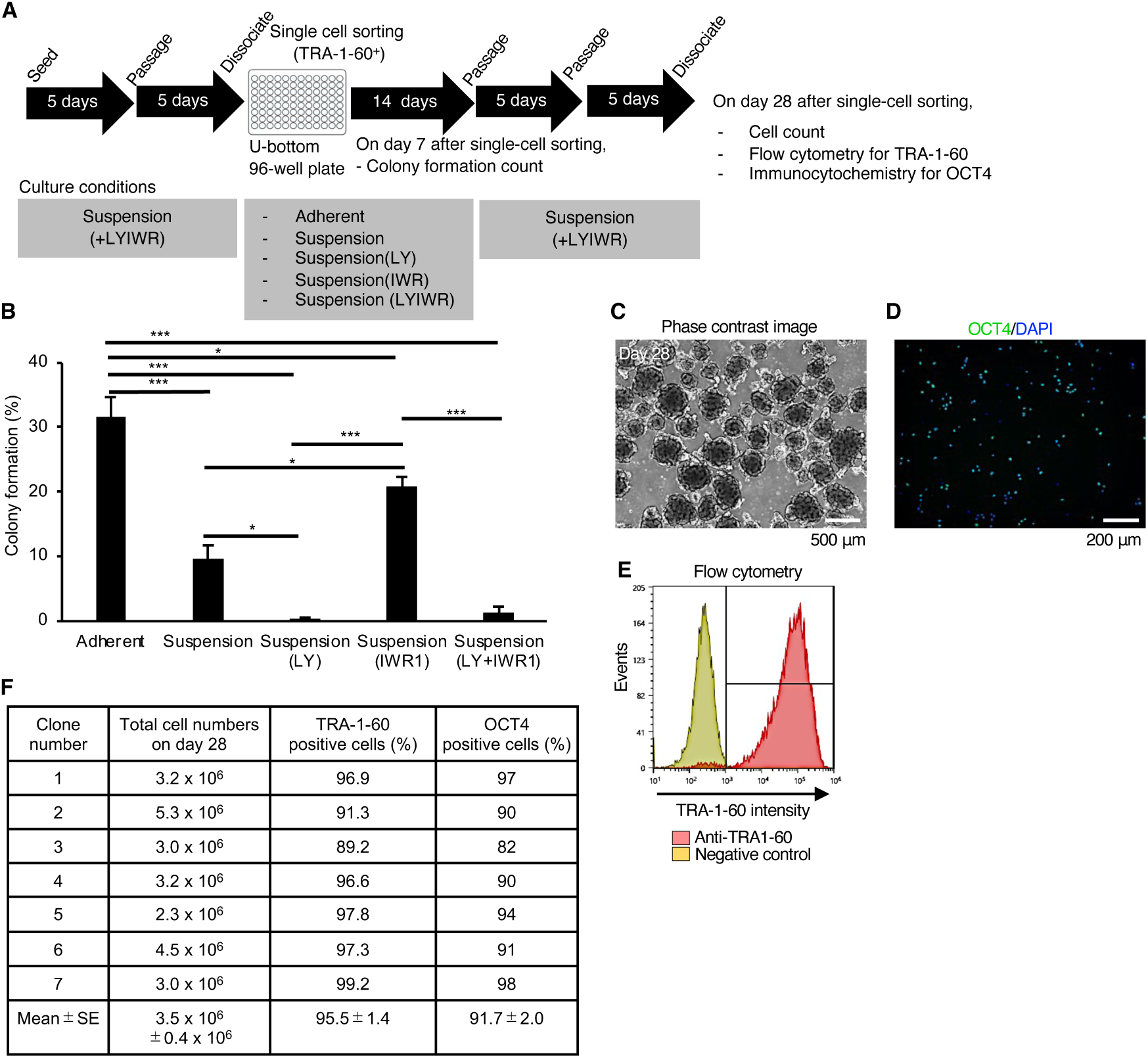
Establishment of single-cell-sorted hiPSC subclones cultured in suspension conditions supplemented with IWR1-endo and LY3333531. (A) Schematics representing the establishment of single-cell-derived hiPSC subclones from 201B7 line. Single-cell-sorted cells were expanded in the culture medium supplemented with IWR-1-endo and LY333531. Formed colonies were picked on day 14, and expanded by repeating passage every 4-5 days under suspension conditions. Characteristic analysis was performed on day 28 after single-cell sorting. (B) On day 7 after single-cell sorting, formed colonies were counted per well in the 96-well plate. The ratio (%) of the colony formation is shown in a bar graph (mean ± SE) (n=3). Statistical analysis was performed using by one-way ANOVA and Tukey’s tests for all graphs. P values < 0.05 were considered statistically significant. *, ** or *** in the graphs indicate P<0.05, P<0.01 or P<0.001, respectively. (C) Phase-contrast images of Clone #2 on day 28 after single-cell sorting. Scale bars: 500 µm. (D) Represented flow cytometry of TRA-1-60 (Clone #2). (E) Represented immunocytochemistry of OCT4 (Clone #2). Scale bars: 200 µm. (F) Summary table of the characterization of single cell-sorted clones. Total cell numbers on day 28, the ratio of TRA-1-60-positive cells (%), and the ratio of OCT4-positive cells (%) are shown (mean ± SE) (n=7).

### Direct freeze and thaw of hiPSCs cultured in suspension conditions

To test the feasibility of our suspension culture method to direct freeze and thaw processes, we froze a hiPSC line, 201B7, in suspension culture conditions using StemFit AK02N medium supplemented with IWR1-endo and LY3333531. Then, we thawed these frozen vials and directly reseeded the cells in suspension conditions supplemented with IWR1-endo and LY3333531 (Figure 8A). By day 10 after reseeding, we expanded the hiPSCs in suspension culture supplemented with IWR1-endo and LY3333531 (Figure 8B). On day 10, we examined three vials for their cell growth and expression of OCT4 and TRA-1-60. The subclones showed more than three million cells and high ratios of OCT4- and TRA1-60-positive cells (Figure 8C-E). We also tested mTeSR1 medium for this process. Three different hiPSC lines, WTC11, 1231A3, and HiPS-NB1RGB, cultured in mTeSR1 supplemented with IWR1-endo and LY3333531 successfully recovered from frozen vials (Figure 8—figure supplement 1A-D). These results indicate that we have successfully frozen and thawed hiPSCs in suspension culture conditions directly.

**Figure 8.**
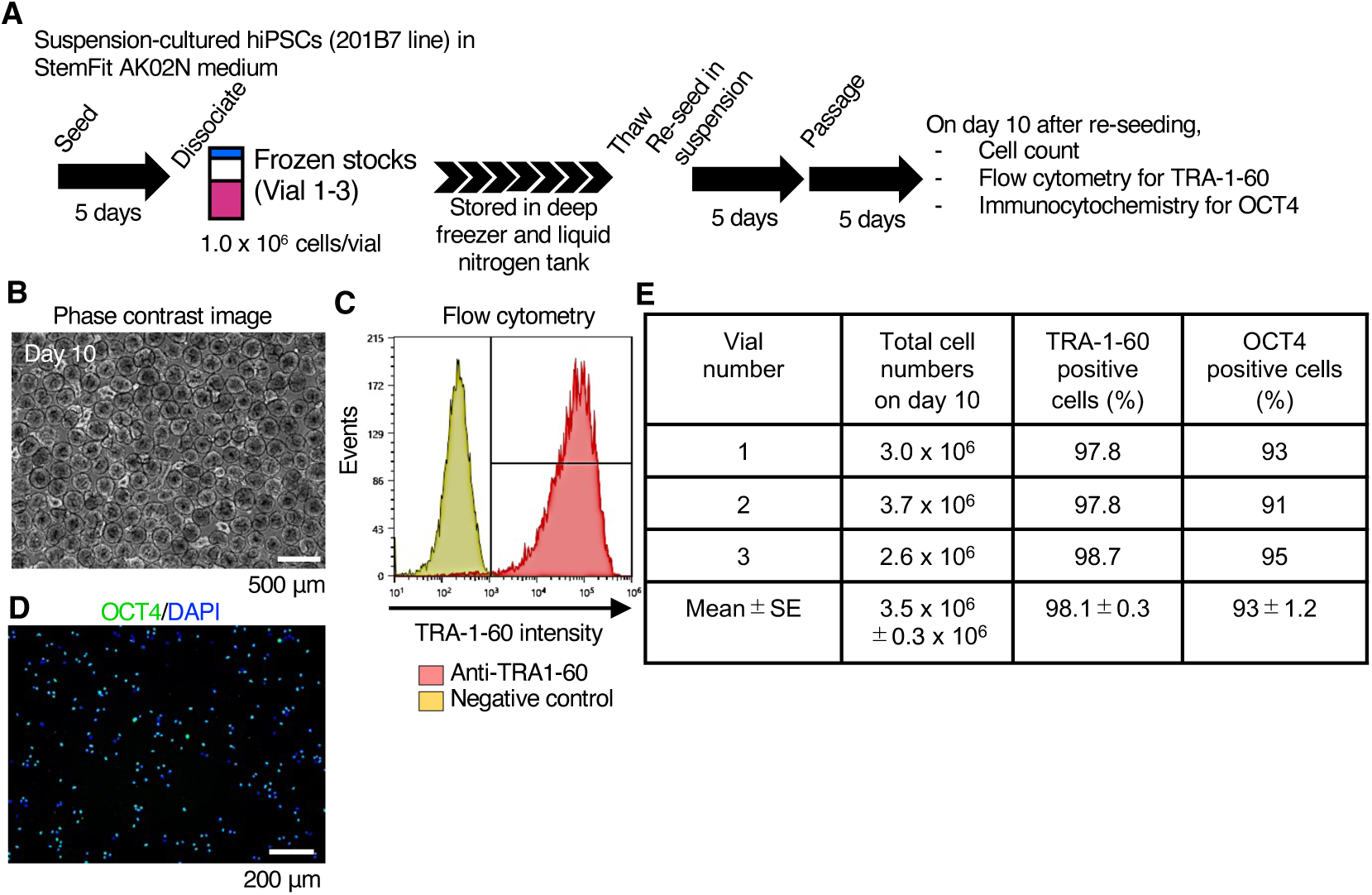
Re-suspension from frozen stocks of hiPSCs under suspension culture conditions supplemented with IWR-1-endo and LY333531. (A) Schematics of re-suspension culture of frozen stocks of single-cells-derived 201B7 clone (Vial 1-3). (B) Represented phase-contrast images on day 10 (Vial 3). Scale bars: 500 µm. (C) Represented flow cytometry of TRA-1-60 (Vial 3). (D) Represented immunocytochemistry of OCT4 (Vial 3). Scale bars: 200 µm. (E) Summary table of the characterization of re-suspension-cultured iPSCs from frozen stocks. Total cell numbers on day 10, the ratio of TRA-1-60-positive cells (%), and the ratio of OCT4-positive cells (%) are shown (mean ± SE) (n=3).

### Establishment of hiPSC lines in complete suspension-culture conditions supplemented with inhibitors of PKCβ and Wnt signaling pathways

For the clinical use of hiPSCs, more secure and efficient establishment of hiPSCs in closed systems is strongly desired. To achieve this, we aimed to establish hiPSCs in suspension culture conditions. Using PBMCs as a starting material, we generated hiPSCs using a novel replication-defective and persistent Sendai virus vector (SeVdp) infection or episomal vector electroporation, as these methods are well-known for producing transgene-free, clinical-grade hiPSCs (Fusaki *et al*, 2009; Nishimura *et al*, 2011) (Figure 9A). PBMCs were infected with SeVdp carrying *OCT4*, *SOX2*, *KLF4*, and *L-MYC* genes. Infected cells were kept cultured by repeating passages every 5 - 6 days. Cell aggregates cultured with IWR1-endo and LY333531 showed uniform spherical structure (Figure 9B), and most of the cells were positive for OCT4 (Figure 9C) and TRA-1-60 (Figure 9D) on day 56. These bulk cells were able to differentiate into three germ layers in an in vitro EB formation assay (Figure 9E) and in teratomas that were transplanted into immunodeficient NOD.Cg-Prkdcscid Il2rgtm1Wjl/SzJ (NSG) mice (Figure 9F). These bulk-hiPSCs maintained normal karyotype (Figure 9G). By single cell sorting with fluorescent labeled TRA-1-60 antibody, we expanded single-cell-derived clones in suspension-culture conditions. A clone (F-10) showed OCT4 and TRA1-60 expressions (Figure 9H-J). In addition, the established clone showed potency to differentiate into derivatives of three germ layers *in vitro* as EBs (Figure 7K) and *in vivo* as teratoma transplanted into NSG mice (Figure. 9L). A normal karyotype was observed in this clone (Figure 9M). SeVdp was nearly extinct in these bulk populations and F-10 clone (Figure 7N). These results demonstrate that, we were successful in establishing transgene-free hiPSC lines using SeVdp infection in suspension-culture conditions in the presence of PKCβ and Wnt inhibitors.

**Figure 9.**
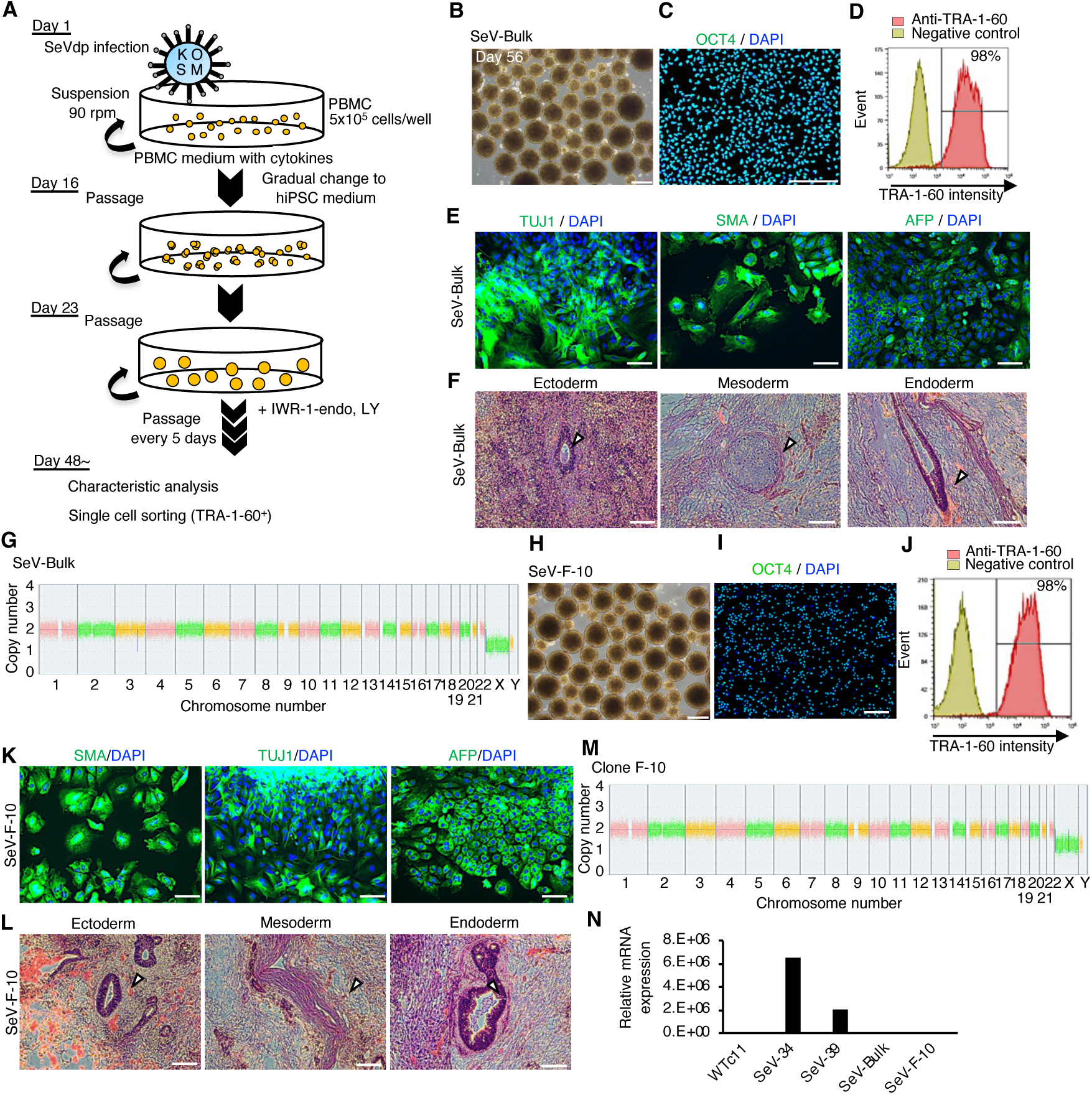
Establishment of hiPSCs in complete suspension conditions using SeVdp. (A) Schematics. (B) Phase-contrast images of PBMCs on day 56 after infection. Scale bars: 400 µm. (C) Immunocytochemistry of OCT4 on bulk-hiPSCs on day 56. Scale bars: 200 µm. (D) Flow cytometry of TRA-1-60 in bulk-hiPSCs on day 61. (E) Immunocytochemistry of TUJ1, SMA, and AFP on differentiated cells in EBs from bulk-hiPSCs on day 56. Scale bars: 100 µm. (F) HE staining of teratoma sections derived from bulk-hiPSCs. White arrowheads indicate representative tissue structures derived from ectoderm, mesoderm, and endoderm. Scale bars: 100 µm. (G) Chromosomal copy numbers detected with CNV array analysis on bulk-hiPSCs. (H) Phase-contrast image of an established hiPSC clone at passage 7. Scale bars: 400 µm. (I) Immunocytochemistry of established clone, F-10, with anti-OCT4 antibody. Scale bars: 200 µm. (J) The percentage of TRA-1-60-positive cells with flow cytometry on established clone F-10 line. (K) HE staining of teratoma sections derived from F-10 clone line. The details are the same as in (F). (L) Immunocytochemistry in EBs from established F-10 clone. Scale bars: 100 µm. (M) Chromosomal copy numbers of F-10 clone line. (N) Residual SeVdp genomic RNA in established hiPSCs with RT-qPCR.

Furthermore, we were successful in establishing single-cell-derived hiPSC lines using the transfection of episomal plasmid vectors in suspension-culture conditions in the presence of PKCβ and Wnt signaling inhibitors. PMBCs were transfected with episomal plasmid vector mix carrying *OCT4*, *SOX2*, *KLF4*, *L-MYC*, *LIN28*, and *mp53DD* and cultured in suspension conditions in the presence or absence of IWR-1-endo and LY33351 with repeated passages to reach enough cell numbers to make cell stocks and characterization (Figure 9—figure supplement 1A). After 36 days of culture, 59% double positive cells for TRA-1-60 and SSEA4 were observed in control, whereas 78% were seen in the presence of IWR-1-endo and LY333531 (Figure 9— figure supplement 1B and C). We performed cell sorting of TRA-1-60 and SSEA4 double positive cells and further repeated 5 more passages in the absence or presence of IWR-1-endo and LY333531. Cell aggregates cultured in culture medium supplemented with IWR-1-endo and LY333531 showed uniform spherical structures (Figure 9—figure supplement 1D), and the most of these cells were TRA-1-60 and SSEA4 double positive (Figure 9—figure supplement 1E). In contrast, cell aggregates cultured without these inhibitors showed lumpy, heterogeneous shapes, and decreased undifferentiated population. Immunostaining of OCT4 on dissociated cell aggregates resulted in higher OCT4 positive cells in IWR-1-endo and LY333531 treated cells (Figure 9—figure supplement 1F and G). We further performed single cell sorting of TRA-1-60-positive cells from bulk-cultured populations at 10 passages (Figure 9—figure supplement 1H and I). We selected 3 clones to be characterized further. These clones showed round spherical shapes in their aggregates and expression of OCT4, NANOG, and TRA-1-60 proteins (Figure 9—figure supplement 1J-L). Pluripotency into three germ layers was confirmed in vitro with EB formation assay (Figure 9—figure supplement 1M) and teratoma formation analysis (Figure. Figure 9—figure supplement 1N). These clones retained a normal karyotype even after a long-culture (Figure 9—figure supplement 1O). These results demonstrate that single-cell-derived hiPSC lines were established using the transfection of episomal plasmid vectors in suspension culture conditions in the presence of PKCβ and Wnt signaling inhibitors.

## Discussion

In this study, we have developed a series of methods to generate and maintain hiPSCs in suspension conditions. First, we have identified compounds that suppress the spontaneous differentiation of hiPSCs in suspension cultures. Based on these findings, we have newly achieved a complete series of culture processes including hiPSC establishment, long-term culture, mass culture, single-cell cloning, and direct freeze and thaw. Our methods are validated in several conventional culture media and many hiPSC lines. Thus, our findings show that suspension culture conditions with Wnt and PKCβ inhibitors (IWRLY suspension conditions) can precisely control cell conditions and are comparable to conventional adhesion cultures regarding cellular function and proliferation. Many previous 3D culture methods intended for mass expansion used hydrogel-based encapsulation or microcarrier-based methods to provide scaffolds and biophysical modulation (Chan *et al*, 2020). These methods are useful in that they enable mass culture while maintaining scaffold dependence. However, the need for special materials and equipment and the labor and cost involved are concerns toward industrial mass culture. On the other hand, our IWRLY suspension conditions do not require special materials such as hydrogels, microcarriers, or dialysis bags, and have the advantage that common bioreactors can be used. Furthermore, we have observed some differences in conventional media used for suspension culture conditions for maintaining self-renewal characteristics, preventing spontaneous differentiation into specific lineages, and performing stability among different experimental times. Overcoming these heterogeneity caused by conventional media, the IWRLY suspension conditions robustly maintain hiPSC self-renewal and pluripotency. Therefore, this IWRLY suspension culture of hiPSCs is advantageous in terms of mass culture, automation, safety assurance, and is expected to have novel industrial applications that could not be achieved by conventional methods.

HiPSCs have been generally considered to be scaffold-dependent and are cultured under adherent monolayer culture conditions (Hayashi & Furue, 2016; Xu *et al*., 2001). On the other hand, some studies reported that floating cultures without external ECM addition or scaffolds have been successfully performed on hiPSCs for their long-term and/or mass expansion (summarized in Table 1). However, dissociation into single cells and amplification from these single cells in suspension culture conditions have never been achieved. Thus, it remains arguable whether culturing human pluripotent stem cells without providing a scaffold is possible. In this study, we demonstrated that many existing hiPSCs quickly acclimatized to the IWRLY suspension culture conditions and that they could be successfully cultured even under harsh conditions such as colony formation from single cells or direct freeze-thaw processes. These findings suggest that hiPSCs can be sufficiently cultured even without scaffolds or exogenous ECM proteins. However, as previous studies have shown, hiPSCs themselves secrete scaffold substances such as ECM proteins, which may affect the status of suspension culture conditions (Kim *et al*., 2019).

On the other hand, it is interesting to see whether and how the properties of hiPSCs cultured in IWRLY suspension culture conditions are altered from the adherent conditions. Our transcriptome results in comparison to adherent conditions show that gene expression associated with cell-to-cell attachment, including E-cadherin (CDH1), is more activated. This may be due to the status that these hiPSCs are more dependent on cell-to-cell adhesion where there is no exogenous cell-to-substrate attachment in the three-dimensional culture. Previous studies have shown that cell-to-cell adhesion by E-cadherin positively regulates the survival, proliferation, and self-renewal of human pluripotent stem cells (Aban *et al*, 2021; Li *et al*, 2012; Ohgushi *et al*, 2010). Furthermore, studies have shown that human pluripotent stem cells can be cultured using an artificial substrate consisting of recombinant E-cadherin protein alone without any ECM proteins (Nagaoka *et al*, 2010). Also, cell-to-cell adhesion through gap junctions regulates the survival and proliferation of human pluripotent stem cells (Wong *et al*, 2006; Wong *et al*, 2004). These findings raise the possibility that the cell-to-cell adhesion, such as E-cadherin and gap junctions, are compensatory activated and support hiPSC self-renewal in situations where there are no exogenous ECM components and its downstream integrin and focal adhesion signals are not forcedly activated in suspension culture conditions. It will be interesting to elucidate these molecular mechanisms related to E-cadherin in the hiPSC survival and self-renewal in IWRLY suspension conditions in the future.

We have identified two compounds as factors that ameliorate the disadvantages of the suspension culture of hiPSCs. Wnt signaling inhibitors specifically inhibited differentiation toward mesendoderm, whereas, the PKCβ inhibitor, LY333531, inhibited ectodermal differentiation and, when combined with IWR-1-endo, maintained the undifferentiated nature of hiPSCs with high efficiency. As for Wnt activation in human pluripotent stem cells, previous studies reported some WNT agonists were expressed in undifferentiated human pluripotent stem cells (Dziedzicka *et al*., 2021; Jiang *et al*, 2013; Konze *et al*, 2014). In suspension culture, cell aggregation causes tight cell-cell interaction. The paracrine effect of WNT agonists in the cell aggregation may strongly affect neighbor cells to induce spontaneous differentiation into mesendodermal cells. Thus, the inhibition of WNT signaling should be effective to suppress the spontaneous differentiation into mesendodermal lineages in suspension culture. Gö6983, a pan-PKC inhibitor, has been used to promote the self-renewal of mammalian PSCs (Dutta *et al*, 2011; Kinehara *et al*, 2013; Rajendran *et al*, 2013; Takashima *et al*, 2014); however, in these cases, the role of specific isoforms in PSC self-renewal and differentiation are not fully elucidated yet. Also, PKC signaling is involved in neural induction conserved in animals shown in sea urchin (Range *et al*, 2013), Xenopus (Otte *et al*, 1988; Otte *et al*, 1989), mouse (Stumpo *et al*, 1995). Our findings suggested the involvement of PKCβ in human neuroectodermal differentiation using hiPSCs. It will be interesting to elucidate the molecular mechanisms of how PKC signaling is involved in neuroectoderm differentiation in the future.

Interestingly, these combinations of chemical inhibitors against WNT and PKC signaling pathways are also being used in the induction and maintenance of naïve human pluripotent stem cells (Bayerl *et al*, 2021; Bredenkamp *et al*, 2019; Guo *et al*, 2017; Khan *et al*, 2021). Previous studies suggested that cell state transition toward naïve state in hPSCs had beneficial effects in the suspension culture of hiPSCs (Lipsitz *et al*., 2018; Rohani *et al*., 2020). While, in suspension-culture, we did not aim to drive hiPSCs to a naïve state by the use of these combinations of chemical inhibitors conditions, we found that naïve pluripotency marker genes were up-regulated in the IWRLY suspension conditions consistently. The question of why the transition to the naïve PSCs type facilitates suspension culture adaptation remains elusive; however, considering early development, the inner cell mass in blastocysts in the earlier stages that maintains pluripotency forms a three-dimensional morphology, whereas epiblasts, where the development is more proceeded, form a flattened epithelial morphology (Sheng, 2015). The same is true for pluripotent stem cells in mice, where naïve mouse embryonic stem cells derived from blastocysts form three-dimensional colonies, whereas epiblast stem cells derived from the later developmental stages form flattened epithelial colonies (Brons *et al*, 2007; Tesar *et al*, 2007). We showed that these differences in the morphology of mouse pluripotent stem cells could be regulated by the activation state of ECM and integrin signaling and that the situation where these signals are not active is suited to the naïve state (Hayashi *et al*, 2007). The same characteristics may be applied to human pluripotent stem cells, although no direct results have yet been shown in human pluripotent stem cells (Hayashi & Furue, 2016). Thus, hiPSCs can be adapted to form three-dimensional colony morphology in IWRLY suspension cultures by shifting their state to a naïve PSCs type, which may facilitate self-renewal in suspension culture.

Last but not least, we successfully generated hiPSC lines from PBMCs in suspension conditions for the first time by adding the abovementioned compounds to the culture medium. Both transduction methods with SeVdp and episomal plasmid vectors enabled the establishment of hiPSCs under suspension conditions in the presence of IWR-1-endo and LY333531. Clonal expansion of hiPSCs was also performed by single-cell sorting with flow cytometry, and the established hiPSCs were successfully characterized for their self-renewal, pluripotency, and genomic integrity. These results indicate that hiPSC lines generated in suspension conditions have the same quality in terms of their pluripotency, self-renewal, and genomic integrity as those generated with conventional adherent conditions. This consistent process from establishing hiPSCs from somatic cells to their mass expansion with precise control of cellular status in suspension culture may pave the way for their stable and automated clinical application toward autologous cell therapy of hiPSCs.

## Methods

### Human subjects

The generation and use of hiPSCs were approved by the Ethics Committee of RIKEN BioResource Research Center, Graduate School of Medicine of Kyoto University, and KANEKA CORPORATION.

### Animal experiments

All animal experiments were approved by the Animal Experimentation Committee of the RIKEN Tsukuba branch and were performed according to the committee’s guiding principles and the “Guide for the Care and Use of Laboratory Animals” published by the National Institute of Health.

### Data availability

All data are available in the main text or the supplementary materials. RNA-seq data are available in the Gene Expression Omnibus as GSE222833. All the data supporting the findings of this study are available from the lead contact upon reasonable request. Source data is provided with this paper. This paper does not report original code.

### HiPSCs cultured in adherent conditions

WTC11 (GM25256 from Coriell Institute) (Hayashi *et al*, 2016), 201B7 (HPS0063 from RIKEN Cell Bank) (Takahashi *et al*., 2007), 454E2 (HPS0077 from RIKEN Cell Bank) (Okita *et al*, 2011), 1383D6 (HPS1006 from RIKEN Cell Bank) (Okita *et al*., 2011), 1231A3 (HPS0381 from RIKEN Cell Bank) (Okita *et al*., 2011), Ff-I14s04 (CiRA foundation, Kyoto University) (Kitano *et al*., 2022), and HiPS-NB1RGB (HPS5067 from RIKEN Cell Bank), which were generated from human neonatal skin fibroblast (RCB0222) (Borisova *et al*, 2022; Shimizu *et al*, 2022), were used as healthy-donor hiPSC lines. With tdTomato-reporter hiPSC lines, we generated knock-in hiPSC lines for PAX6-TEZ (constructed from 454E2 line) and SOX17-TEZ (constructed from 1383D6 line) using CRISPR-Cas9 genome editing (Tsukamoto *et al*, 2021) (Figure. S1A). These hiPSCs were cultured in StemFit AK02N medium (Cat#AK02N, Ajinomoto, Co., Ltd., Tokyo, Japan). Medium change was performed every day and passaged at 80%–90% confluency after 6-7 days of culture. At passage, PBS/EDTA solution (0.5 mM, Cat#06894-14, Nacalai Tesque) was used to dissociate hiPSC colonies, and these cells were seeded at a density of 2,500 cells/cm^2^. 10 µM Y-27632 (Cat#HY-10071, FUJIFILM Wako Pure Chemical Corporation, Osaka, Japan) and 0.25 µg/cm^2^ iMatrix-511 silk (Cat#892021, Matrixome Inc, Osaka, Japan) were added to the culture dish on seeding day. These hiPSC cells were cultured in a CO_2_ incubator (Forma Steri-Cycle i160, Thermo Fisher Scientific) with a gas conditions at 5% CO_2_, 21% O_2_, >95% humidity, and 37 °C.

### Suspension-culture of hiPSCs

Suspension culture with rotation at 90 rpm was performed with a plate shaker (Cat#WB-101SRC, WAKENBTECH, Kyoto, Japan or #0081704-000, TAITEC, Tokyo, Japan) installed in a CO_2_ incubator (Cat#Steri-Cycle i160, Thermo Fisher Scientific, MA, USA) and operated under high humidity continuously during the whole culture period. To start the culture hiPSCs in suspension conditions, 4 × 10^5^ cells were seeded in one well of a low-attachment 6-well plate (Cat#MS-80060, Sumitomo Bakelite, Tokyo, Japan) with 4 mL of StemFit AK02N medium, StemScale PSC suspension medium (A4965001, Thermo Fisher Scientific, MA, USA), or mTeSR medium (Stem Cell Technologies, Vancouver, Canada) supplemented with 10 µM Y-27632. This plate was placed onto the plate shaker in the CO_2_ incubator (Forma Steri-Cycle i160, Thermo Fisher Scientific). The medium without Y-27632 was changed every day unless otherwise specified. On days 3 - 5, the hiPSC aggregates were dissociated with Accutase (Cat#12679-54, Nacalai Tesque) at 37°C for 10 min. The dissociated cells were counted with an automatic cell counter (Model R1, Olympus) with Trypan Blue staining to detect live/dead cells. These cell suspension was spun down at 200 rpm for 3 minutes, and the supernatant was aspirated. The cell pellet was re-suspended with a new culture medium at an appropriate cell concentration and used for the next suspension culture. This passage was performed every 5 days unless otherwise specified. To screen for factors that inhibit spontaneous differentiation of hiPSCs, chemicals or recombinant proteins were added to the culture medium (listed in Supplementary Table S1). These hiPSC cells were cultured in a CO_2_ incubator with a gas conditions at 5% CO_2_, 21% O_2_, >95% humidity, and 37 °C.

### Bioreactor culture of hiPSCs

A frozen stock of hiPSCs was pre-cultured twice in adherent conditions to prepare enough cell numbers. To prepare enough hiPSCs to start large-scale culture, hiPSCs were pre-cultured in iMatrix-511MG (Cat#892 005, Matrixome, Inc, Osaka, Japan) or Vitronectin (VTN-N) Recombinant Human Protein, Truncated (Cat#A14700, Thermo Fisher Scientific, MA, USA)- coated cell culture flasks with StemFitAK03N (Cat#AK03N, Ajinomoto, Co., Ltd., Tokyo, Japan) including 20 µM IWR-1-endo and 1 µM LY333531. A 30-mL stirred suspension bioreactor (BWV-S03A, Able Co., Tokyo, Japan) was used according to a previous study (Matsumoto *et al*., 2022). The medium change was manually performed every other days. After 3-4 days of culture, the formed hiPSC aggregates were dissociated with TrypLE SELECT (Cat#A12859-01, Thermo Fisher Scientific, MA, USA). As for large-scale hiPSCs culture, the reactor system BioFlo320 (Eppendorf, Hamburg, Germany) was used according to a previous study (Kropp *et al*., 2016). Perfusion-culture was started with 3.2 - 4.8 × 10^7^ cells in 320 mL of StemFit AK03N medium with IWR-1-endo and LY333531. To maintain the lactate concentration below a certain level and to regulate the pH, the culture was carried out by increasing the amount of medium perfusion per unit time in accordance with the cell proliferation transition. To prevent pH decrease, CO_2_ concentration was regulated by feedback control in the reactor system. After 3-4 days of culture, the formed hiPSC aggregates were dissociated with TrypLE SELECT (Cat#A12859-01, Thermo Fisher Scientific, MA, USA) collected for making cell stocks (∼300 tubes). This perfusion culture was repeated three times (P1, P2, and P3) and the cells were prepared at each expansion step.

### HiPSC generation in suspension culture conditions

HiPSCs were generated from healthy donor-derived PBMCs (Precision for Medicine). Thawed PMBCs from a vial containing around 1 × 10^7^ cells were pre-cultured in one well of low attachment 6-well plate including 4 mL of StemSpan-AOF (Cat#ST100-0130, STEMCELL Technologies, Vancouver, Canada) supplemented with recombinant human IL-6 (100 ng/mL), IL-3 (10 ng/mL), SCF (300 ng/mL), TPO (300 ng/mL), and FLT3 ligand (300 ng/mL) (all from FUJIFILM Wako Pure Chemical Corporation, Tokyo Japan). After 24 h of incubation with continuous stirring at 37°C/5% CO_2_/21% O_2_, PBMCs were spun down with centrifugation at 200 × g for 10 min at low deceleration speed and resuspended in StemSpan ACF for cell counting. For episomal vetor transduction, 2.5 × 10^6^ cells of PMBCs were centrifuged at 200 × g for 10 min with low deceleration speed and electroporated using Nucleofector 2b device (Lonza, Basel, Switzerland) with iPS cell Generation Episomal vector Mix (Takara Bio, Shiga, Japan) and Amaxa Human CD34^+^ Cell Nucleofector kit (Lonza, Basel, Switzerland) according to manufacturer’s protocol. 5 × 10^5^ of electroporated cells were seeded in one well of a low attachment 6-well plate in 4 mL StemSpan ACF with the cytokines mentioned above. Suspension-culture was performed with continuous agitation at 90 rpm. Stem span ACF medium were gradually replaced with StemFit AK02N medium. Formed cell aggregates were passaged with Accutase on day 16 in the presence of 10 µM Y27632, and suspension-culture was continued until the cell numbers reached a sufficient amount for characterization. 10 µM IWR-1-endo and 1 µM LY333531 were added from day 3.

SeVdp infection was performed with Sendai Reprogramming Kit (CytoTune EX-iPS of virus solution, ID Pharma, Tsukuba, Japan) according to the manufacturer’s protocol with some modifications. Briefly, pre-cultured 1 × 10^6^ PMBCs were centrifuged at 200 × g for 10 min, with low deceleration speed, and resuspended in 2 mL of StemSpan ACF with cytokines. PMBCs were gently mixed with 2 mL of virus solution prepared at MOI = 5 per 1 × 10^6^ cells. 5 × 10^5^ infected cells were seeded in one well of low-attachment 6-well plate at a total volume of 4 mL with StemSpan ACF plus cytokines. Suspension-culture was initiated with continuous agitation at 90 rpm. Stem span ACFs were gradually replaced with StemFit AK02N as mentioned above, and the cell aggregates were passaged with Accutase on day 16 in the presence of 10 µM Y-27632. Suspension-culture and passages continued until cell reached a sufficient number for characterization. 10 µM IWR-1-endo and 1 µM LY333531 were added from day 23. These cells were cultured in a CO_2_ incubator (Forma Steri-Cycle i160, Thermo Fisher Scientific) with a gas conditions at 5% CO_2_, 21% O_2_, >95% humidity, and 37 °C.

### Three germ layer differentiation *in vitro*

For embryoid body (EB) formation assay, suspension-cultured hiPSC lines were dissociated with Accutase, and 1.0 x 10^4^ cells were seeded in each well of an EZ-BindShut 96-well-V plate (Cat#4420-800SP AGC TECHNO GLASS CO., LTD, Shizuoka, Japan) with 100 µL Stem Fit AK02N medium supplemented with 10 µM Y-27632. Before culture, the 96-well-V plate was centrifuged at 200 x g for 3 min for efficient cell mass formation. The next day, the culture medium was switched to DMEM high Glucose (Cat#08458-16, Nacalai tesque, Kyoto, Japan) supplemented with 10% fetal bovine serum (here after referred as EB medium). On day 8, EBs were transferred into 0.1% (w/v) Gelatin Solution (Cat#190–15805, FUJIFILM Wako Pure Chemical Corporation, Tokyo Japan) -coated 12-well plate and further cultured in EB medium for another 8 days. The medium was changed every day. EBs were fixed with 4% paraformaldehyde in Dulbecco’s Phosphate Buffered Saline (D-PBS) (Cat#14249-24, Nacalai tesque, Kyoto, Japan) for 10 min at room temperature and used for immunostaining. The differentiation was validated by immunostaining against each germ layer markers.

The differentiation potency of suspension-cultured hiPSCs toward three germ layers was also evaluated by culturing in germ layer specific differentiation medium. As in the maintenance conditions, 4 × 10^5^ hiPSC were seeded in one well of a low-attachment 6-well plate with 4 mL of StemFit AK02N medium supplemented with 10 µM Y-27632. This plate was placed onto the plate shaker in the CO_2_ incubator. Next day, the medium was changed to the germ layer specific differentiation medium. For ectodermal differentiation, suspension-cultured hiPSCs spheroids were cultured with Stem Fit AK02N without C medium supplemented with 10 µM SB431542 (Cat#198-16543, FUJIFILM Wako Pure Chemical Corporation, Tokyo, Japan) and 10 µM DMH1 (Cat#041-33881FUJIFILM Wako Pure Chemical Corporation, Tokyo, Japan) for 7 days. For mesodermal differentiation, suspension-cultured hiPSCs spheroids were cultured with RPMI1640 medium (FUJIFILM Wako Pure Chemical Corporation, Tokyo, Japan) supplemented with B-27 supplement, minus insulin (A1895601, Thermo Fisher Scientific, MO, USA) and 7.5 µM CHIR99021 (Cat#034-23103, FUJIFILM Wako Pure Chemical Corporation, Tokyo, Japan) for 1 day, and continuously with RPMI1640 medium supplemented with B-27 supplement for 2 days. For endodermal differentiation, suspension-cultured hiPSCs spheroids were cultured with RPMI1640 medium supplemented with 1 mM Sodium pyruvate (FUJIFILM Wako Pure Chemical Corporation, Tokyo, Japan), 1 x NEAA (FUJIFILM Wako Pure Chemical Corporation, Tokyo, Japan), 80 ng/mL Activin A (R&D Systems, MN, USA), 55 µM 2-mercaptethanol (FUJIFILM Wako Pure Chemical Corporation, Tokyo, Japan), 50 ng/mL FGF2 (R&D Systems, MN, USA), 20 ng/mL BMP4, and 3 µM CHIR99021 for 2 days, and continuously with RPMI1640 medium supplemented with 1 mM Sodium pyruvate, 1× NEAA, 80 ng/mL Activin A, 55 µM 2-mercaptethanol, and 0.5% Knockout serum replacement (KSR; Thermo Fisher Scientific, MO, USA) for 2 days.

### Neuroectoderm and endoderm differentiation of reporter hiPSC lines

Reporter hiPSC lines, PAX6-TEZ and SOX17-TEZ were used as positive controls for in vitro neuroectoderm and endoderm differentiation, respectively. PAX6-TEZ was seeded in each well of a 24 well-plate coated with 0.25 µg/cm^2^ iMatrix-511 silk with 1 mL StemFit AK02N medium supplemented with 10 µM Y-27632. Next day, the culture medium was switched to Stem Fit AK02N medium without supplement C, instead of containing 10 µM SB431542 and 10 µM DMH1. SOX17-TEZ was seeded in each well of a 24 well-plate (same as above), and on the next day, the culture medium was switched to StemFit AK02N medium without supplement C instead of containing 3 µM CHIR99021 and 10 ng/mL Activin A (Cat#014-23961, FUJIFILM Wako Pure Chemical Corporation, Tokyo, Japan). The medium was changed every day. On day 7, tdTomato expression was observed under all-in-one fluorescent microscope (BZ-X800; KEYENCE).

### Cardiomyocyte differentiation

Cardiomyocyte differentiation was performed according to a modified method described previously (Funakoshi *et al*, 2016). Both large-scale suspension cultured hiPSCs and typical adherent-cultured hiPSC were collected by centrifugation after dissociation into single cell with TrypLE Select. The cells were suspended in 1.5 mL cardiomyocyte differentiation media (CDM), consisting of StemPro34 medium (Thermo Fisher Scientific) supplemented with 2 mM Glutamax (Thermo Fisher Scientific), 50 μg/mL ascorbic acid, 4 × 10^-4^ M monothioglycerol (Sigma-Aldrich), 150 μg/mL transferrin (Roche), and with Matrigel (Cat#354277, Corning, NY, USA)), 10 μM Y-27632 and 2 ng/mL human recombinant BMP4 (R&D Systems) and then cultured in ultra-low attachment 6 well plate (Corning) in 5%CO_2_/5% O_2_. After 24 h, 1.5 mL CDM with 6 ng/mL human recombinant activin A (R&D Systems) and 5 ng/mL bFGF (R&D Systems) were added into the wells. After 3 days, medium was changed to 3 mL CDM with 10 ng/mL VEGF (R&D Systems), SB431542 (Sigma-Aldrich), Dorsomorphin (Sigma-Aldrich) and 1 µM IWP-3 (Stemgent). After 7 days, medium was changed to 3 mL CDM with 10 ng/mL VEGF (R&D Systems). The medium was changed every 2 days. The cells were collected at 15 days after differentiation and analyzed by flow cytometry.

### Dopaminergic progenitor cells differentiation

Dopaminergic progenitor cells differentiation was performed according to a modified method described previously (Doi *et al*, 2020). Both large-scale suspension cultured hiPSCs and typical adherent-cultured hiPSC were collected by centrifugation after dissociation into single cell with TrypLE Select. The cells were suspended in 1 mL dopaminergic progenitor cells differentiation medium-1 (DPM-1), consisting of Glasgow’s minimum essential medium (Thermo Fisher) supplemented with 8% knockout serum replacement (Thermo Fisher Scientific), 1% MEM Non-Essential Amino Acids Solution (Thermo Fisher Scientific), 1 mM sodium pyruvate (Sigma-Aldrich), 0.1 mM 2-mercaptoethanol (FujiFilm Wako) and 100 nM LDN193189 (Stemgent), and were then seeded in iMatrix-coated 24-well plates at a density of 1 × 10^6^ cells/well with 10 µM Y-27632 and 500 nM A-83-01 (FujiFilm Wako). After 1 and 2 days, medium was changed to DPM-1 with 500 nM A-83-01, 100 ng/mL recombinant human FGF8 (FujiFilm Wako), and 2 μM purmorphamine (FujiFilm Wako). After 3 to 6 days, medium was changed every day to DPM-1 with 500 nM A-83-01, 100 ng/mL recombinant human FGF8, 2 μM purmorphamine, and 3 μM CHIR99021 (FujiFilm Wako). After 7 to 11 days, medium was changed every day to DPM-1 with 3 μM CHIR99021. On differentiation day 12, cells were dissociated using TrypLE select and suspended in the dopaminergic progenitor cells differentiation medium-2 (DPM-2), consisting Neurobasal medium (Thermo Fisher Scientific), 2% B27 supplement (without Vitamin A, Thermo Fisher), 1% GlutaMAX (Thermo Fisher Scientific), 10 ng /mL human recombinant glial cell-derived neurotrophic factor (FujiFilm Wako), 200 mM ascorbic acid, 20 ng / mL human recombinant brain-derived neurotrophic factor (FujiFilm Wako), and 400 μM dibutyryl cAMP (FujiFilm Wako), and were then plated on U-shaped 96-well plates (Thermo Fisher Scientific) at a density of 2 × 10^4^ cells/150 µL/well with 10 µM Y-27632. After 15 to 26 days, half of medium was changed every 2 days to DPM-2. Dopaminergic progenitor cells differentiation efficiency was analyzed by flow cytometry at 12 days after differentiation. On differentiation day 12, cells were labeled with anti-CORIN antibody (Clone 5B6, Sigma-Aldrich) and secondary antibody (A11001, Thermo Fisher).

### Hepatocyte differentiation

Hepatocyte differentiation was performed according to a modified method described previously (Si-Tayeb *et al*, 2010). Both large-scale suspension-cultured and typical adherent-cultured hiPSCs were collected by centrifugation after dissociation into single cell with TrypLE Select. These hiPSCs were cultured on Matrigel-coated 6-well plates until 80% confluency with StemFit medium (Ajinomoto). To initiate differentiation, the medium was changed to 1 mL Hepatocyte differentiation medium (HDM) consisting of RPMI1640 medium (Thermo Fisher) supplemented with 1× GlutaMax (Thermo Fisher) and 1× B27 Supplement minus vitamin A, with the addition of 100 ng/mL Activin A (R&D Systems). After 1 to 3 days, medium was changed every day to HDM with 100 ng/mL Activin A. After 4 to 8 days, medium was changed every day to HDM with 20 ng/mL human recombinant BMP-4 (R&D Systems) and 20 ng/mL human recombinant FGF-4 (R&D Systems). After 9 to 13 days, medium was changed every day to HDM with 20 ng/mL human recombinant HGF (R&D Systems). After 14 to 24 days, medium was changed every 2 days to Hepatocyte Culture Medium (HCM; Lonza) with 20 ng/mL human recombinant Oncostatin M (R&D Systems). On differentiation day 25, the albumin concentration in the culture supernatant was determined by ELISA using an anti-human albumin antibody (Betyl Laboratories, Catalog # E88-129).

### Teratoma formation

Suspension-cultured hiPSCs were dissociated with Accutase and then resuspended with 1 mL of StemFit AK02N medium supplemented with 10 µM Y-27632. 1 x 10^6^ cells were collected by centrifugation at 200 × g for 3 m and suspended in ice-cold 50% Matrigel solution: StemFit AK02N with 10 µM Y-27632 = 1:1). Cells were injected into testis or leg of NOD.Cg-Prkdcscid Il2rgtm1Wjl/SzJ (NSG) mice (The Jackson Laboratory, Bar Harbor, ME, USA) using an 18-G needle. Two to three months later, teratomas were collected and fixed with 4% paraformaldehyde in D-PBS. Paraffin-embedded sectioning and hematoxylin-eosin (HE) staining was performed. Three germ layer derivatives were observed under CKX53 microscope with a DP22 camera and CellSens software (EVIDENT, Tokyo, Japan).

### Apoptosis assay of suspension-cultured iPSCs

Suspension-cultured hiPSCs were dissociated with Accutase. The dissociated cells were aliquoted into 1 x10^5^ cells/100 µL with a binding buffer consisted of 10 mM HEPES, 140 mM NaCl, 2.5 mM CaCl_2_, then added with 5 µL Annexin V (Alexa Fluor 680) conjugates (Cat# A35109, Thermo Fisher scientific) and 1 µL DAPI solution (Cat# 340-07971, FUJIFILM Wako, Pure Chemical Corporation, Tokyo, Japan). After incubation for 15 minutes at room temperature in dark conditions, 400 µL of cold binding buffer was added. The apoptotic cells were immediately analyzed with a flow cytometer (SH800S, SONY, Tokyo, Japan). As a positive control of apoptosis, suspension-cultured hiPSCs were treated with Staurosporine (1 µM) (Cat#S1421, Selleck Biotech, Tokyo, Japan), one of the apoptosis-inducing factors, for 2 hours before collection.

### Karyotyping

For virtual karyotyping, genomic DNA (gDNA) was extracted from hiPSCs using a DNeasy Blood & Tissue Kit (Cat#69504, QIAGEN, Hulsterweg, Netherland) and was used for microarray assay. Virtual karyotyping was performed with GeneChip Scanner System 3000 using Karyostat Assay arrays (Cat#905403, both from Thermo Fisher Scientific, MA, US) according to the manufacturer’s protocol. Data were analyzed using the Chromosome Analysis Suite (ChAS) and Affymetrix GeneChip Command Console software programs.

G-band analysis was performed using the common Giemsa staining method with hiPSCs fixed by Carnoy’s fixtative (3:1 ratio of methanol: glacial acetic acid).

### qRT-PCR

Total RNA was extracted with a FastGene RNA premium kit (Cat#FG-81250, NIPPON Genetics, Tokyo, Japan) and used for reverse transcription reaction. cDNA was synthesized by using a ReverTra Ace qPCR RT kit (Cat#FSQ-101, TOYOBO, Osaka, Japan) with random primers. Real-Time qPCR reactions were performed with a QuantStudio 3 System (Thermo Fisher Scientific) using THUNDERBIRD Probe qPCR Mix (Cat#QPS-101, TOYOBO) with TaqMan probes (listed in Supplementary Table 2) (Thermo Fisher Scientific) according to the manufacturer’s instructions. Gene expression was described as the fold change relative to the control sample value (ΔΔCt method) after normalization to the corresponding GAPDH or β-Actin values, unless otherwise specified. For the residual SeV detection, data were normalized to the expression of GAPDH and displayed as a relative fold increase to hiPSC line established with episomal vector (WTC11 line). SeVdp-infected fibroblasts on passage 1 (SeV-34 and -39) were used as positive controls.

### Immunocytochemistry

Immunocytochemistry was performed on adherent cells. Suspension-cultured hiPSCs were dissociated with Accutase and transiently cultured as adherent in iMatrix-coated culture dishes for 3 h before immunocytochemistry. The cells were fixed with 4% paraformaldehyde in D-PBS for 10 min at room temperature, then permeabilized in D-PBS containing 0.1% Triton X-100 for another 10 min. Cells were incubated with primary antibodies in D-PBS containing 0.1% Bovine Serum Albumin (BSA; Cat#017-22231, FUJIFILM Wako Pure Chemical Corporation, Tokyo, Japan) overnight at 4 °C. The secondary antibodies were incubated for 1 hour at room temperature in D-PBS containing 0.1% BSA. Fluoro-KEEPER Antifade Reagent, Non-Hardening Type with DAPI (Cat#12745-74, Nacalai tesque, Kyoto, Japan) was used for nuclear counterstaining. Fluorescence images were taken with an all-in-one fluorescent microscope (BZ-X800; KEYENCE, Osaka, Japan). The primary and secondary antibodies used in this study are listed in Supplementary Tables 3 and 4, respectively.

### Automatic capillary Western blot (Simple Western Assays)

To detect the expression of PAX6 or SOX17, adherent- or suspension-cultured PAX6-TEZ or SOX17-TEZ hiPSCs were collected on day 10 (Passage 2). The cell lysate was prepared from 1 × 10^6^ cells with sodium dodecyl sulfate-polyacrylamide gel electrophoresis (SDS-PAGE) sample buffer solution without 2-ME (2x) (Cat#30567-12, Nacalai Tesque) supplemented with 100 mM Dithiothreitol (DTT) (Cat#14130-41, Nacalai Tesque). To detect phosphorylated PKCβ, cell lysates were collected on day 5 of suspension-cultured hiPSCs with extraction buffer mentioned above, containing 1% phosphatase inhibitor cocktail (Cat#07574-61, Nacalai Tesque). Samples were denatured at 95°C for 5 min. Western blotting was performed with a capillary automatic western blotting device (Simple Western, Wes; Bio-Techne, CA, USA). Jes/Wes 12- to 230-kDa separation module for Wes, 8 × 25 capillary cartridges (Cat#SM-W004, ProteinSimple), and Anti-Rabbit/Goat/Mouse Detection Module kit (Cat#DM-002/DM-001/DM/006, ProteinSimple) were used. Preparation of reagents and sample loading were conducted according to the manufacturer’s instructions. The data were analyzed and quantified by compass for simple Western software program (ProteinSimple). GAPDH was used as the reference for normalization. The primary antibodies used in this study are listed in Supplementary Table 3.

### Flow cytometry

Flow cytometry of self-renewal markers, TRA-1-60 and SSEA4 was performed as described in our previous study. Briefly, adherent- or suspension-cultured hiPSCs were dissociated with 0.5 mM EDTA solution in D-PBS or Accutase, respectively. Dissociated cells (0.5 or 1 × 10^6^ cells) were collected and centrifuged at 200 × g for 3 minutes and resuspended in 500 or 1000 µL PBS containing 0.1% BSA and 0.5 mM EDTA, then incubated with or without anti-TRA-60 antibody or anti-SSEA4 antibody for 1 hour at 4 °C under oblique light conditions. After washing with PBS, cells were resuspended in 500 mL D-PBS containing 0.1% BSA and 0.5 mM EDTA, and filtrated with 35-mm cell strainer (Cat#352235, FALCON, Thermo Fisher Scientific). For PAX6-TEZ and SOX17-TEZ hiPSC lines, adherent- or suspension-cultured hiPSCs were dissociated into single cells and collected cells were resuspended in 500 mL PBS containing 0.1% BSA and 0.5 mM EDTA. After passing through the 35-mm cell strainer, cells were analyzed for tdTomato. The primary and secondary antibodies used in this study are listed in Supplementary Table 3 and 4, respectively. Flow cytometry was performed with SH800 cell Sorter (SONY, Tokyo, Japan).

### RNA-Seq

Total RNA was extracted using the FastGene RNA premium kit, and strand-specific library preparation was performed. The prepared library was sequenced using a NovaSeq6000 (Illumina, Inc, CA, USA). Sequencing was performed in a 150 bp × 2 paired-end conFigureuration with a data output of about 6 Gb per sample (∼ 20 million paired reads). Library preparation and sequencing was performed in GENEWIZ (Azenta, MA, USA). To identify differentially regulated genes, sequencing data was analyzed with a CLC Genomics Workbench (QIAGEN, Hulsterweg, the Neterlands) and R package, edgeR (v3.30.3) in R language (v 4.1.0). An MA plot (log2 fold change versus mean average expression) comparing transcriptomes between suspension- and adherent-culture from RNA-seq data. Transcripts with log2 fold change ≧ 2 or ≦ -2 (FDR <0.01) are highlighted with red and blue dots, respectively. The extracted genes were analyzed for gene ontology enrichment and biological pathways using the R package, clusterProfiler (v3.16.1). GSEA was also performed using clusterProfiler. An enrichment map was used to visualize the GSEA results with enrichment score (ES) and false discovery rate (FDR) values. Software used for the analysis is listed in Supplementary Table 5.

### Statistical analysis

Statistical analysis was performed using Student’s t-test, Dunnett’s test for multiple comparisons with a single control condition, and one-way ANOVA and Tukey’s tests for multiple comparisons with all the conditions. P values < 0.05 were considered statistically significant. *, ** or *** in the graphs indicate P<0.05, P<0.01 or P<0.001, respectively. No statistical methods were used to predetermine sample size. The experiments were not randomized and the investigators were not blinded to allocation during experiments and outcome assessment.

## Acknowledgments

We thank Dr. Bruce Conklin for providing us with the WTC11 hiPSC line; Dr. Shinya Yamanaka, Dr. Kazutoshi Takahashi, Dr. Masato Nakagawa, and Dr. Keisuke Okita for providing us with 201B7, 454E2, 1231A3, 1383D6, and Ff-I14s04 hiPSC lines. We would like to express our sincere gratitude to Dr. Kaoru Saijo, Mr. Daiki Kondo and Mr. Masahiko Yamada for their technical assistance, and Ms. Kumiko Omori for her administrative support. This work was funded by a grant from AMED (23bm1423010h0001) to Y.Ha., a collaborative research grant from Kaneka corporation to Y.Ha., and an endowment to CiRA foundation. We would like to thank Editage [http://www.editage.com] for editing and reviewing this manuscript for English language.

## Author Contributions

Conceptualization: YHA, Methodology: MMT, SK, CW, KT, MI, NN YHA, Investigation: MMT, SK, YHE, TW, TS, YA, KT, MI, MN, MU, TMK, MN, TK, RM, MS, NN, KN, YN, YHA, Visualization: MMT, SK, TS, YH, Funding acquisition: MMT, TN, YK, YHA, Project administration: MMT, NN, TN, YK, YHA, Supervision: MMT, TN, YN, NT, YK, YHA, Writing – original draft: MMT, KS, MT, YHA, Writing – review & editing: MMT, YHA

## Competing Interest Statement

YHA received collaborative research grant from KANEKA corporation during this study. SK, KT, MI, YK, TN, TK, RM, MS, and NN were employed with KANEKA corporation during this study. MMT, SK, KT, MI, YK, and YHA. are inventors of patents arising from this work (WO/2021/162090 and PCT/JP2020/005255). The other authors declare no competing interests.

---

**Figure 1—figure supplement 1.**
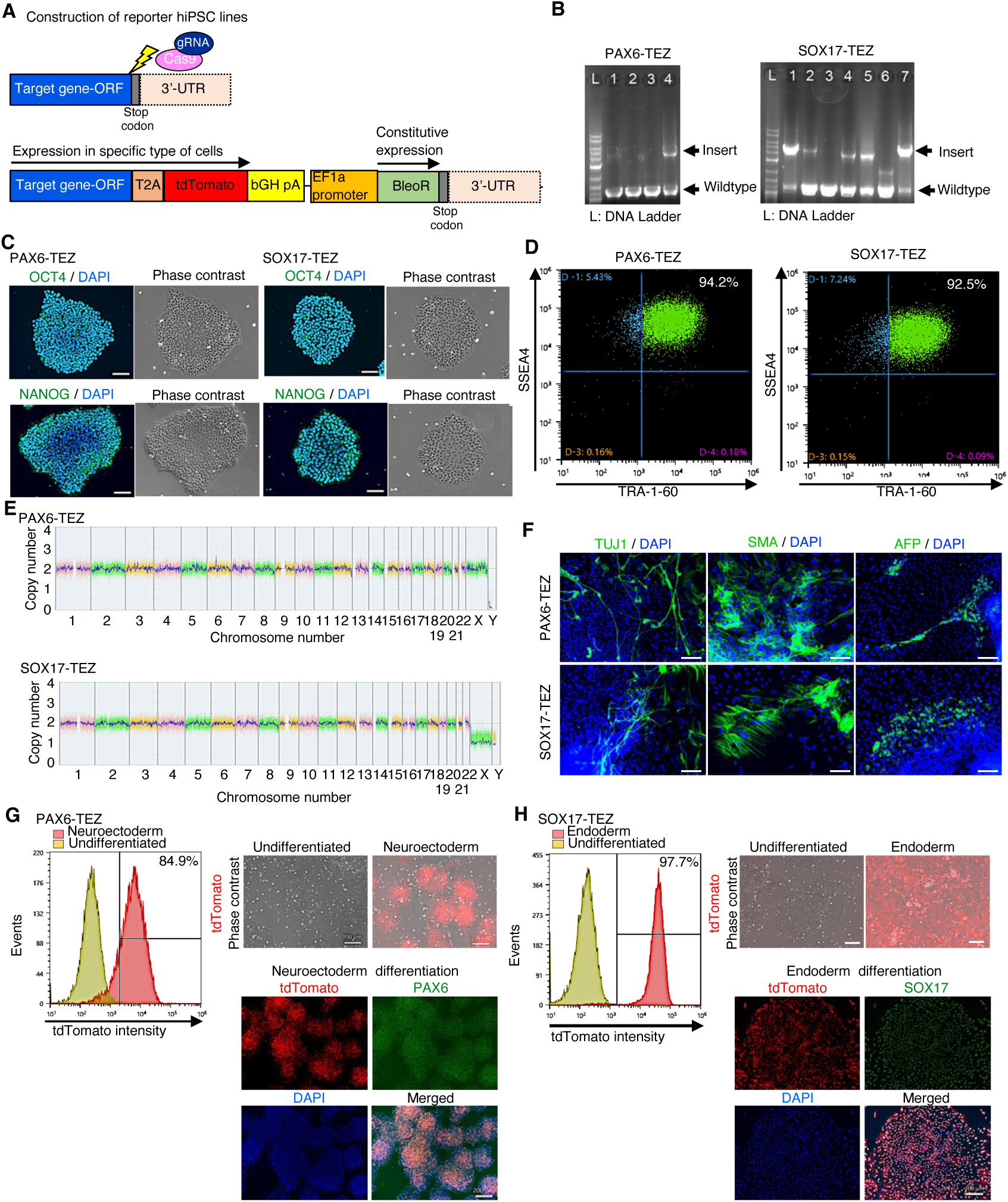
Characterization of PAX6-tdTomato and SOX17-tdTomato reporter lines. (**A**) Schematic of construction of reporter hiPSC lines. CRISPR/Cas9 was employed to knock-in the 2A-tdTomato gene and EF1a promoter-driven Zeocin (Bleomycin) resistance gene sequence (TEZ) to the 3’ terminal of SOX17 or PAX6 open reading frame (ORF), respectively. (B) Genotyping of selected clones by genomic PCR. Arrows indicate alleles of wildtype or insertion in the PAX6 (left panel) or SOX17 (right panel) genes. (C) Phase-contrast images of representative colonies and immunocytochemistry of pluripotency markers: OCT4 and NANOG. Left panels: PAX6-TEZ. Right panels: SOX17-TEZ. (D) Flow cytometry of cell surface markers: TRA-1-60 and SSEA4. Left: PAX6-TEZ. Right: SOX17-TEZ. (E) Chromosomal copy numbers detected with CNV array analysis of PAX6-TEZ (top) and SOX17-TEZ (bottom). (F) Immunocytochemistry of differentiated cells in EBs from established bulk-hiPSCs on day 56. Anti-TUJ1, -SMA and -AFP antibodies were used. Scale bars: 100 µm. (G) Differentiation potency of PAX6-TEZ in the neural ectoderm. Left: Flow cytometry of tdTomato using PAX6-TEZ cultured in neuroectodermal differentiation medium for 7 days. Right upper: Overlay images of phase-contrast and tdTomato fluorescence of PAX6-TEZ cultured in neuroectodermal differentiation medium (upper) or maintenance medium (lower). Scale bars: 200 µm. Right lower: Images of tdTomato fluorescence and endogenous PAX6 protein detected with immunocytochemistry with anti-PAX6 antibody. Nuclear staining was performed with DAPI. Scale bars: 200 µm. (H) Differentiation potency of SOX17-TEZ in the endoderm. Left: Flow cytometry of tdTomato using SOX17-TEZ cultured in endodermal differentiation medium for 7 days. Right upper: Overlay images of phase-contrast and tdTomato fluorescence of SOX17-TEZ cultured in endoderm differentiation medium (upper) or maintenance medium (lower). Right lower: Images of tdTomato and endogenous SOX17 protein detected by immunocytochemistry with anti-SOX17 antibody. Nuclear staining was performed with DAPI. Scale bars: 200 µm.

**Figure 2—figure supplement 1.**
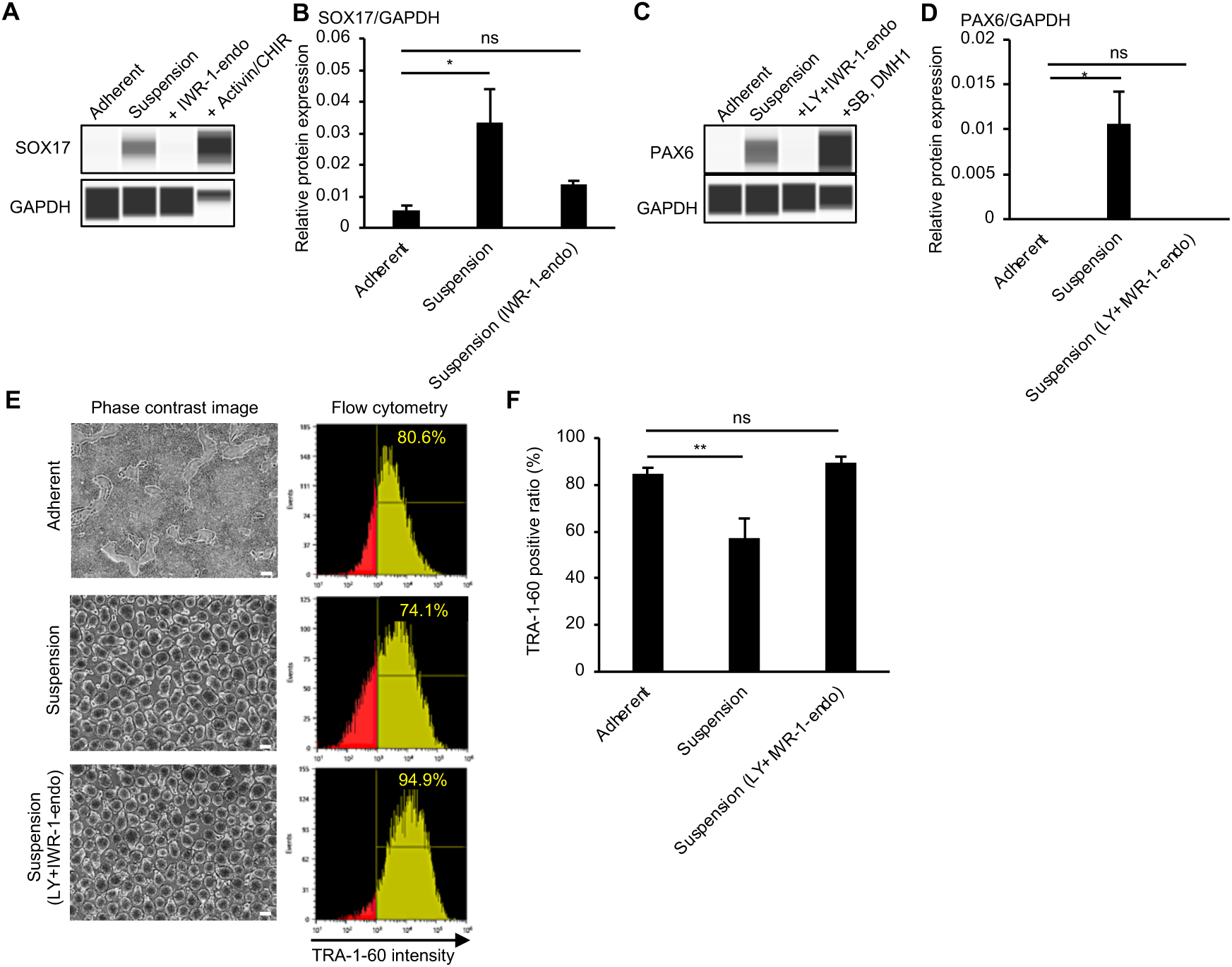
Effects of PKCβ and IWR-1-endo on suspension-cultured hiPSCs at the protein level. (A) Simple western blotting of adherent- or suspension-cultured hiPSCs (SOX17-TEZ) in the presence or absence of IWR-1-endo. SOX17-TEZ cultured in endoderm differentiation medium (Actinin A + CHIR) was used as a positive control for SOX17 expression. (B) Quantification of SOX17 expression. The protein expression was normalized to GAPDH. Data are presented as mean ± SE (n=3). (C) Simple western blotting of adherent- or suspension-cultured hiPSCs (PAX6-TEZ) in the presence or absence of IWR-1-endo and LY333531. PAX6-TEZ cultured in neurectoderm differentiation medium (SB431542+DMH1) was used as the positive control for PAX6 expression. (D) Quantification of expression of PAX6. The protein expression was normalized to GAPDH. Data are presented as mean ± SE (n=3). (E) Phase-contrast images and representative flow cytometry data of pluripotency marker, TRA-1-60, in adherent- or suspension-cultured hiPSCs (1383D6 line) on day10 (Passage 2). Suspension-culture was performed in the presence or absence of IWR-1-end and LY333531. The percentage of TRA-1-60 positive cells is shown in the right corner. (F) Quantified data for TRA-1-60 positive ratio (%). The percentage of TRA-1-60 positive cells is shown as mean ± SE (n=3).

**Figure 2—figure supplement 2.**
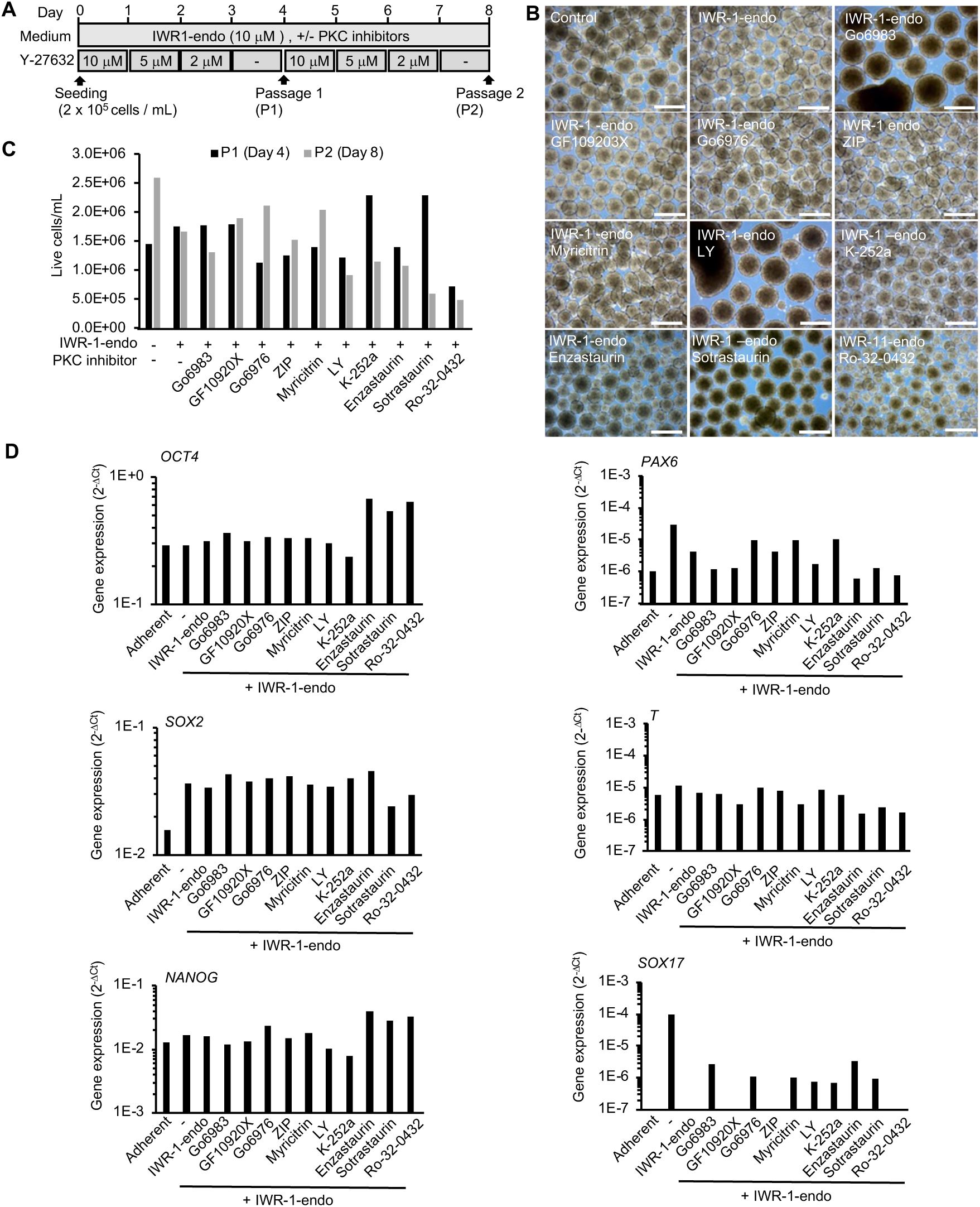
Activity of different type of PKC inhibitors on spontaneous differentiation of suspension-cultured hiPSCs. (A) Schematic representing suspension culture of hiPSCs supplemented with PKC inhibitors. Healthy donor-derived hiPSCs (201B7 line) were grown under continuous agitation in the presence of Y-27632, IWR-1-endo and PKC inhibitors. Passage was performed every four days. (B) Phase-contrast images of suspension cultured hiPSCs on day 8 (P2) in the absence or presence of IWR-1-endo and each PKC inhibitors. Scale bars, 500 µm. (C) Cell growth of hiPSCs (201B7). Graph shows live cell numbers counted on day 4 and day 8 in suspension-culture (n=1). The PKC inhibitors added are labelled in the graphs. (D) Gene expression in hiPSCs cultured in adherent or suspension conditions with each PKC inhibitor. RT-qPCR was performed on cDNAs prepared from day 8 (passage 2) samples (n=1). qPCR was performed for pluripotency (OCT4, SOX2, NANOG) and early differentiation markers (PAX6, T, SOX17). Gene expression was normalized to beta-ACTIN.

**Figure 3—figure supplement 1.**
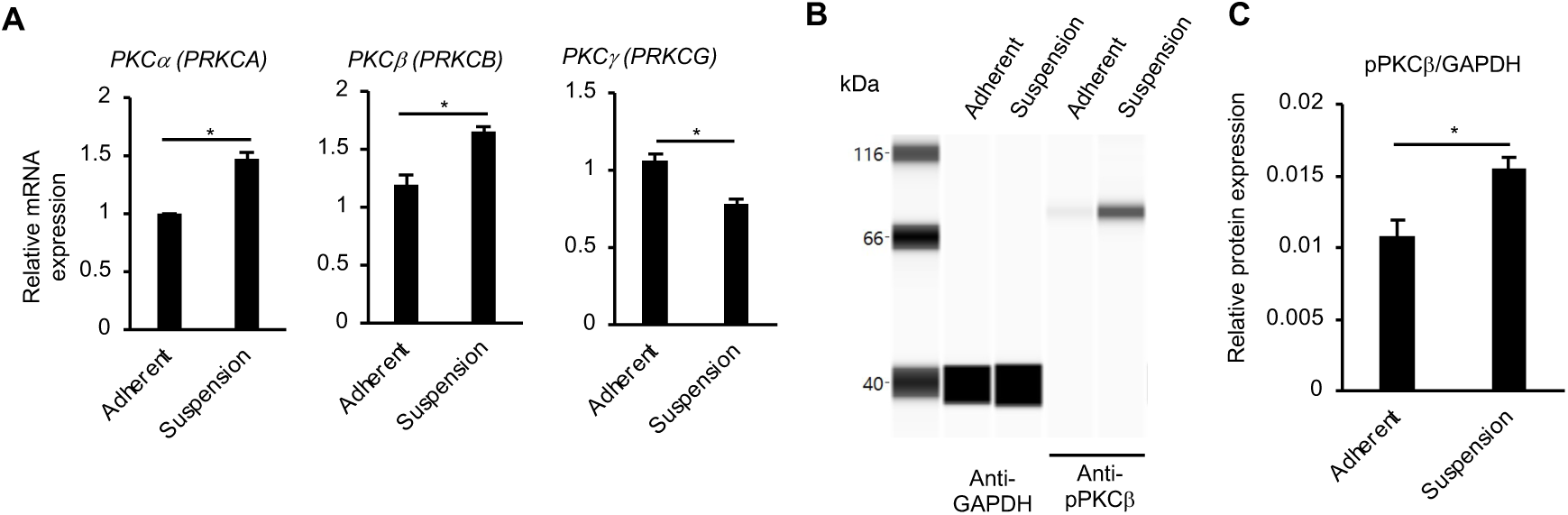
Up-regulation of PKC genes in suspension-cultured hiPSCs. (A) RNA expression of conventional PKC isoform genes in hiPSCs as detected by RT-qPCR. cDNA was prepared from adherent- or suspension-cultured hiPSCs on day 5. The gene expression was normalized to GAPDH. Results are displayed as relative fold increase to adherent-culture. Data are presented as mean ± SE (n=3). P-values were statistically analyzed by Student’s t-test. (B) Detection of phosphorylated PKCβ protein (pPKCβ) in suspension cultured hiPSCs on day 5 using an automated capillary western blot (Simple Western) assay. Adherent-cultured hiPSCs were used as a control. (C) Quantification of pPKCβ expression. The protein expression was normalized to GAPDH. Data are shown as mean ± SE (n=3). P-value was statistically analyzed using Student’s t-test. Scale bars, 200 µm.

**Figure 3—figure supplement 2.**
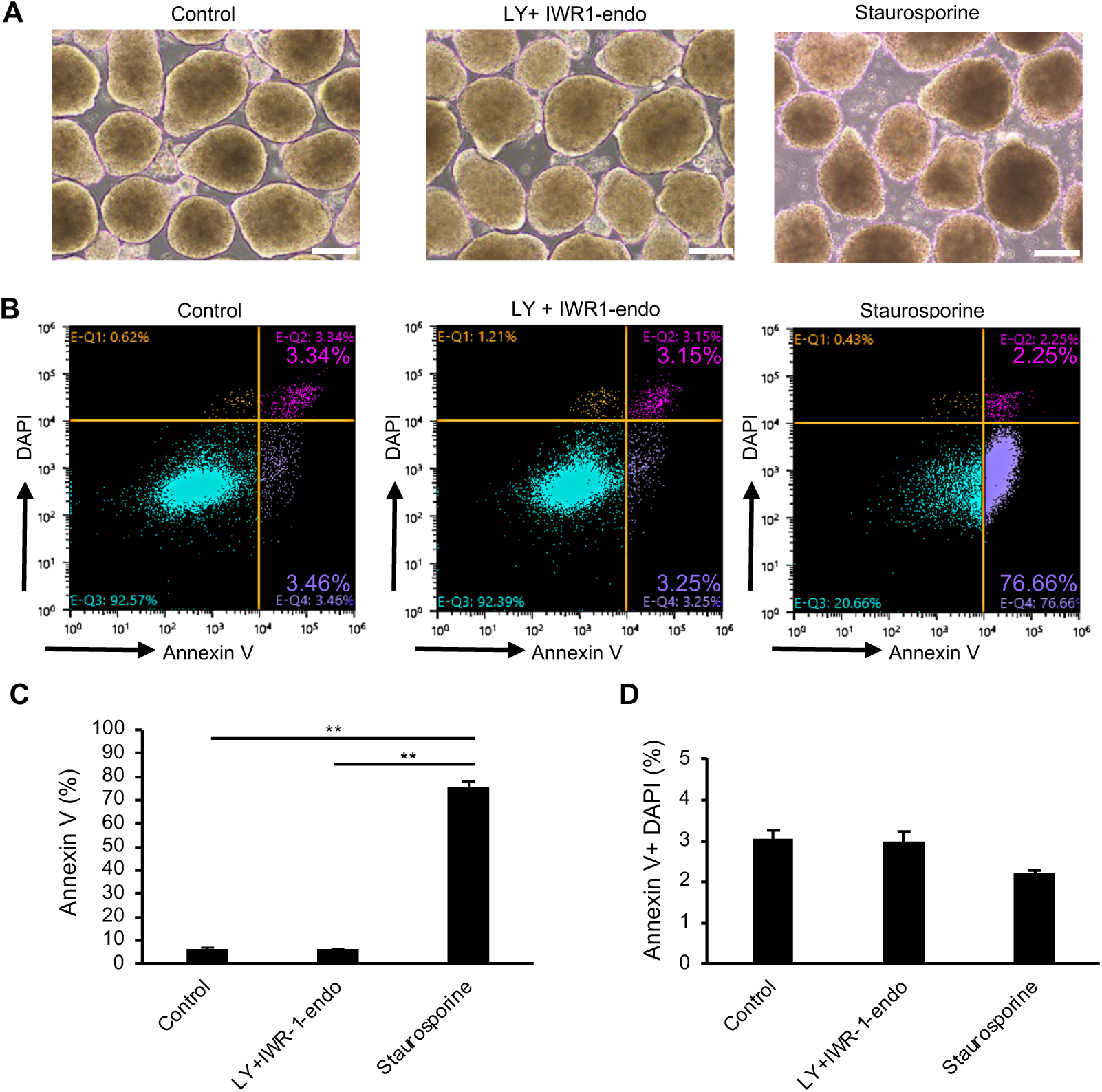
Cell death of suspension cultured iPSCs after the treatment with LY333531 and IWR-1-endo. (A) Phase-contrast images of suspension-cultured hiPSCs (WTC11 line) on day 10 (Passage 2). Cells were culture in StemFit AK02N medium with no supplement (left), with LY333531 + IWR-1-endo (middle) for 10 days, with Staurosporine for 2 hours. Scale bars: 200 µm. (B) Evaluation of apoptotic cells by flow cytometry using Annexin V (Alexa Fluor 680) conjugates. (C) Bar-graph indicating Annexin V-positive cells (%) in each culture conditions. (D) Bar-graph indicating double positive cells for Annexin V and DAPI (%) in each culture conditions. Data are presented as mean ± SE (n=3). Statistical analysis was performed using by one-way ANOVA and Tukey’s tests for all graphs. P values < 0.05 were considered statistically significant. ** in the graphs indicate P<0.01.

**Figure 4—figure supplement 1.**
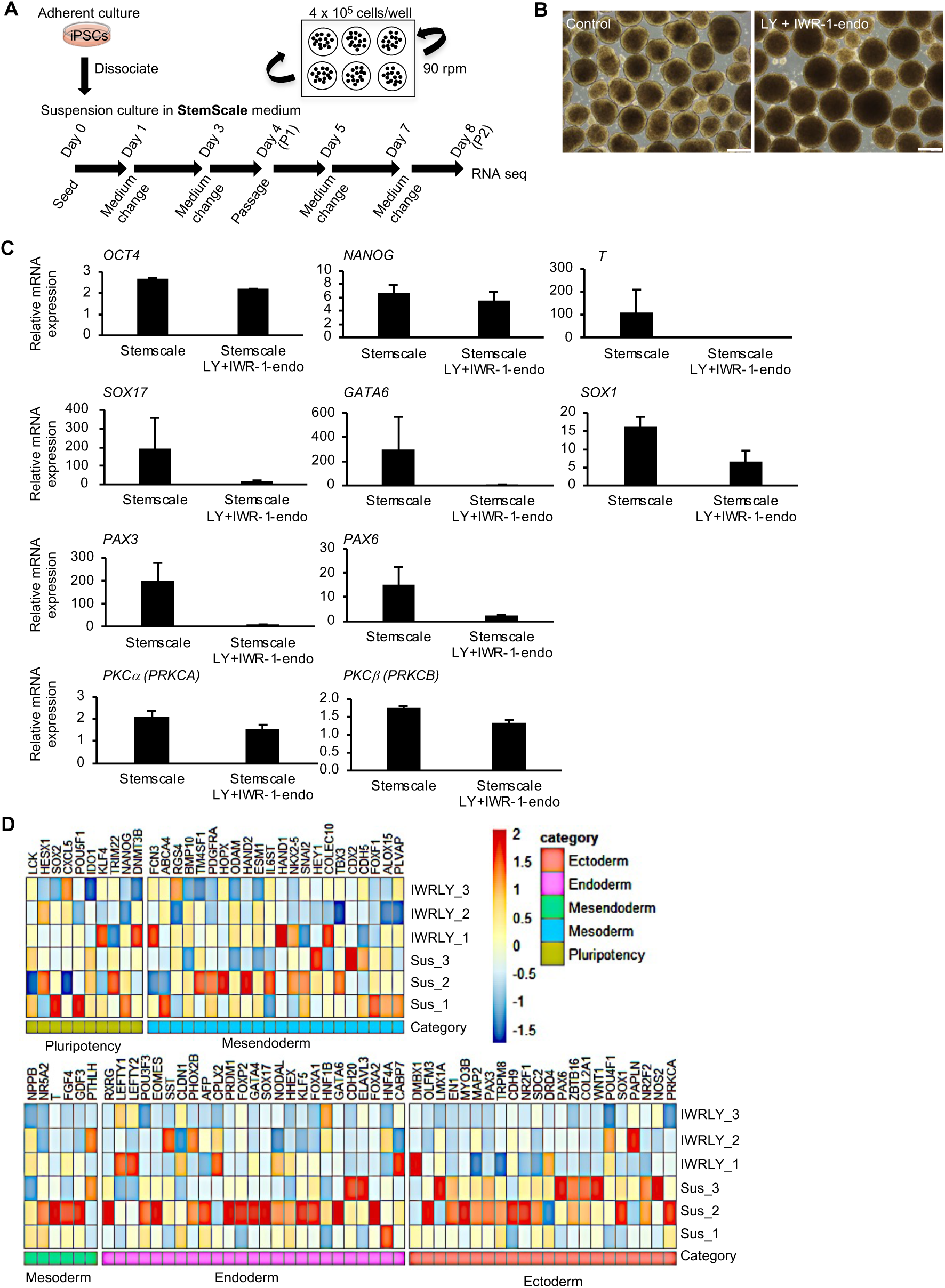
Generality of the inhibitory effects of IWR-1-end and LY333531 on spontaneous differentiation of hiPSCs cultured in suspension condition. (A) Schematic representing suspension culture of hiPSCs using StemScale medium (Thermo Fisher Scientific) as the basic culture medium. hiPSCs were grown under continuous agitation (90 rpm) in the presence or absence of IWR-1-endo and LY333531. Passage was performed every four days according to the manufacture’s protocol. cDNAs were prepared from day 8 (P2) and used for RNA seq. (B) Phase-contrast images of suspension cultured hiPSCs on day 8 in the presence or absence of IWR-1-endo and LY333531. Scale bars, 400 µm. (C) RNA expression of pluripotency and early differentiated genes in hiPSCs, as detected by RT-qPCR. cDNA was prepared from suspension-cultured hiPSCs on day 8 in the presence or absence of IWR-1-endo and LY33353. The gene expression was normalized to GAPDH. Data are presented as mean ± SE (n=3). (D) Heat map of marker genes for pluripotency and differentiation to three germ layers from RNA-seq data in suspension-cultured iPSCs without compounds (Sus_1, Sus_2 and Sus_3) or with IWR-1-endo and LY333531 (IWRLY_1, IWRLY_2 and IWRLY_3). The listed genes were classified as pluripotent, mesendoderm, mesoderm, endoderm, and ectoderm, as defined in the Scorecard Assay (Thermo Fisher Scientific).

**Figure 4—figure supplement 2.**
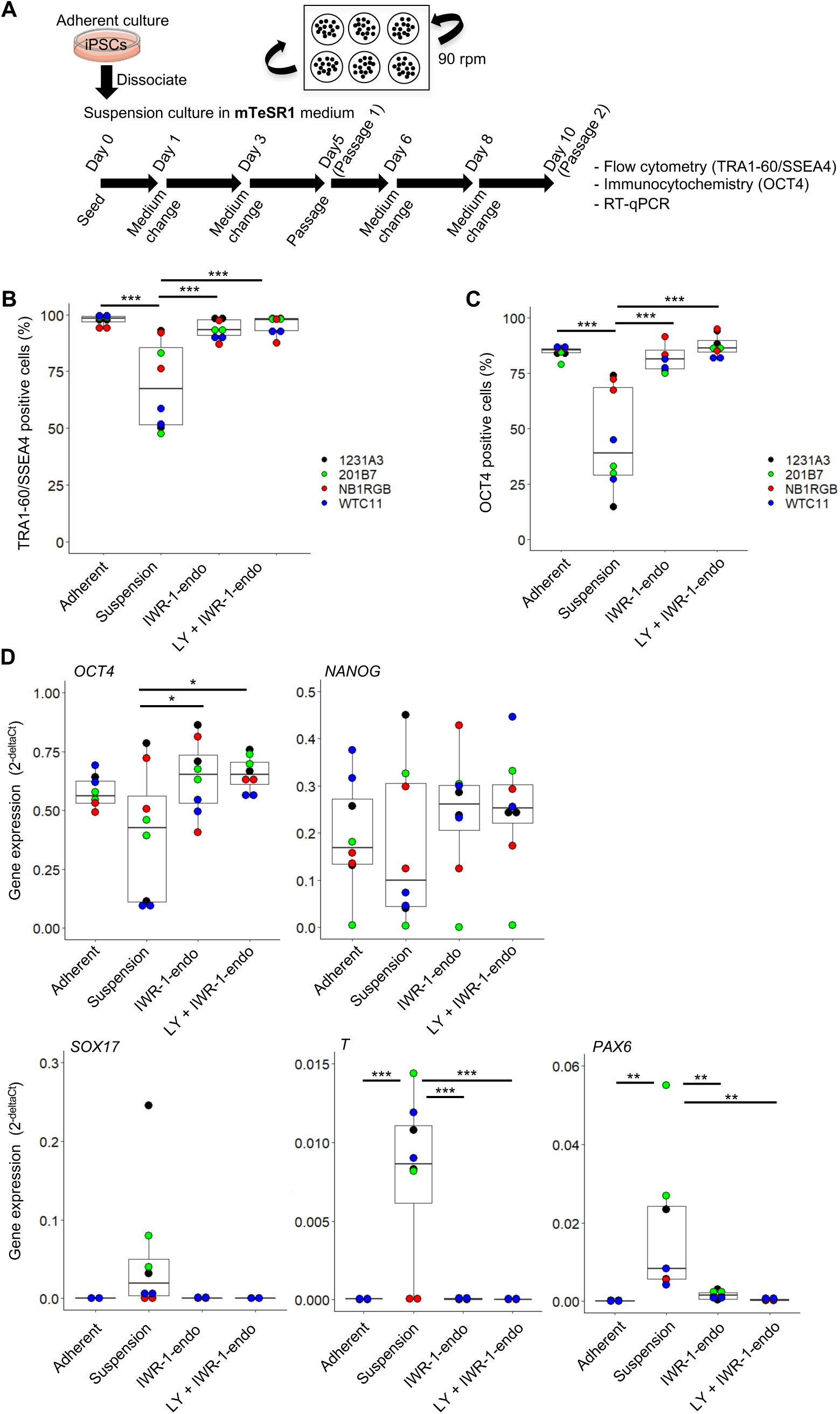
Suspension culture of multiple hiPSC lines in mTeSR1 medium supplemented with or without IWR-1-endo and LY333531. (A) Schematics of suspension culture of WTC11, 1231A3, HiPS-NB1RGB (NB1RGB) and 201B7 hiPSC lines. After suspension culture of these 4 hiPSC lines in mTeSR1 medium with or without IWR-1-endo and LY333531, characteristic analysis was performed on day10 (Passage 2). Suspension culture was performed twice in each cell lines (n=2 each from 4 cell lines). (B) Box plots of flow cytometry for TRA-1-60 and SSEA4 double positive cells (%). Used cell lines are indicated by the different colored circles shown on the right side of the graph. (C) Box plots of OCT4 positive cells (%). (D) Box plots of RT-qPCR. Differentiation markers (*SOX17, T,* and *PAX6*), undifferentiated markers (*OCT4* and *NANOG*), and naïve pluripotency markers (*KLF2* and *KLF5*) were assessed. Statistical analysis was performed using by one-way Anova and Tukey’s test for all graphs. P values < 0.05 were considered statistically significant. *, ** or *** in the graphs indicate P<0.05, P<0.01 or P<0.001, respectively.

**Figure 5—figure supplement 1.**
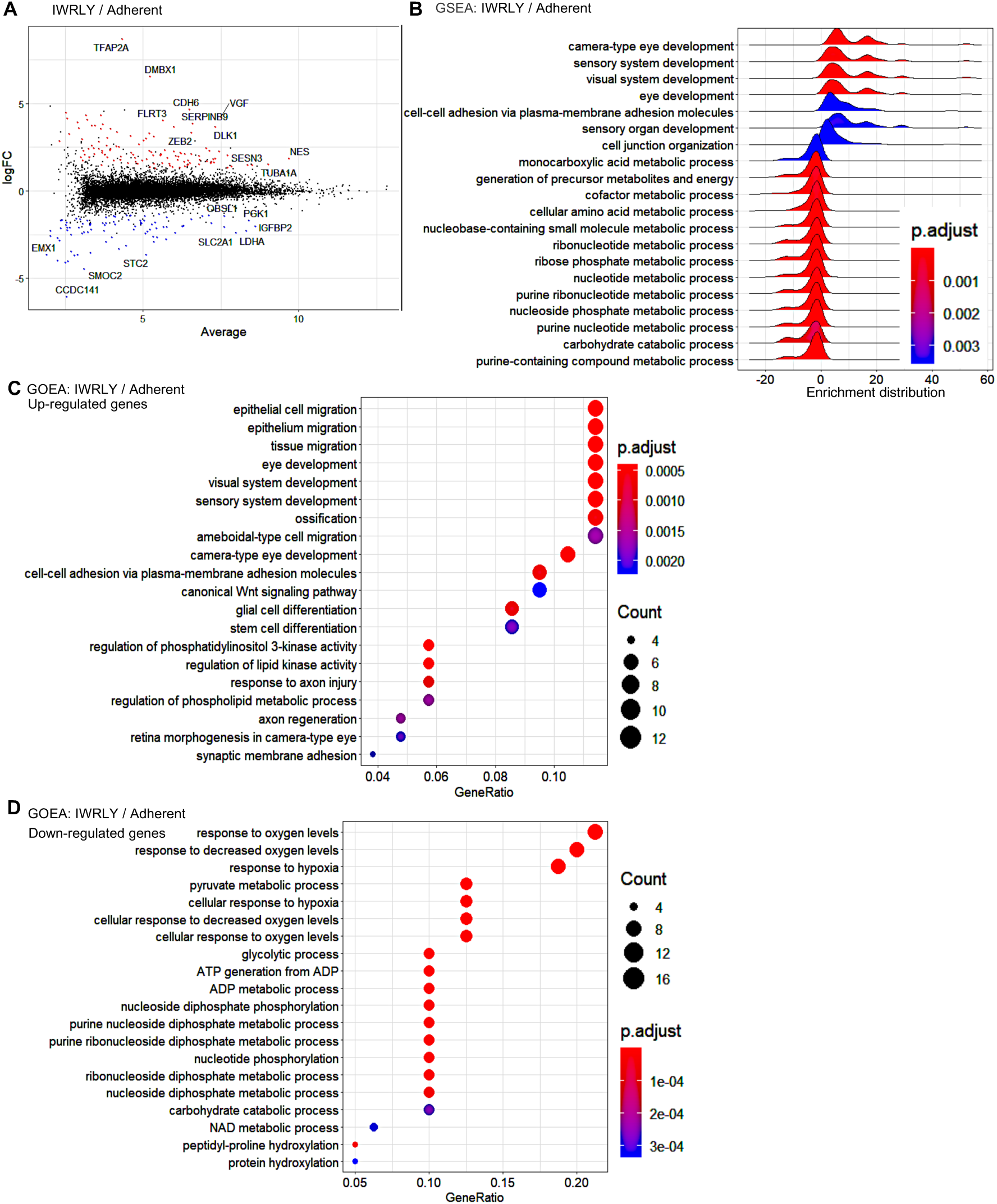
Global gene expression of suspension-cultured hiPSCs with IWR-1-endo and LY333531 in comparison to adherent-cultured hiPSCs. (A) MA plot (log2 fold change versus mean average expression) comparing the transcriptomes between adherent- and suspension-cultured hiPSCs treated with IWR-1-endo and LY33353 (IWRLY). Transcripts with log2 fold change ≧ 2 or ≦ -2 (FDR <0.01) are highlighted with red or blue dots, respectively. The names of the representative genes are shown in the plot. (B) GSEA of the gene sets of suspension-cultured hiPSCs with IWRLY compared to adherent-cultured hiPSCs. Analysis was performed on both up-regulated and down-regulated genes. Statistically significant enrichment is shown. Adjusted P-values are illustrated from blue to red as low to high. (C and D) GOEA of the gene sets of suspension-cultured hiPSCs with IWRLY compared to adherent-cultured hiPSCs. Analysis was performed on up-regulated genes in (C) and down-regulated genes in (D), respectively. Results are ranked by significance (adjusted P values) and/or counted gene numbers.

**Figure 5-figure supplement 2.**
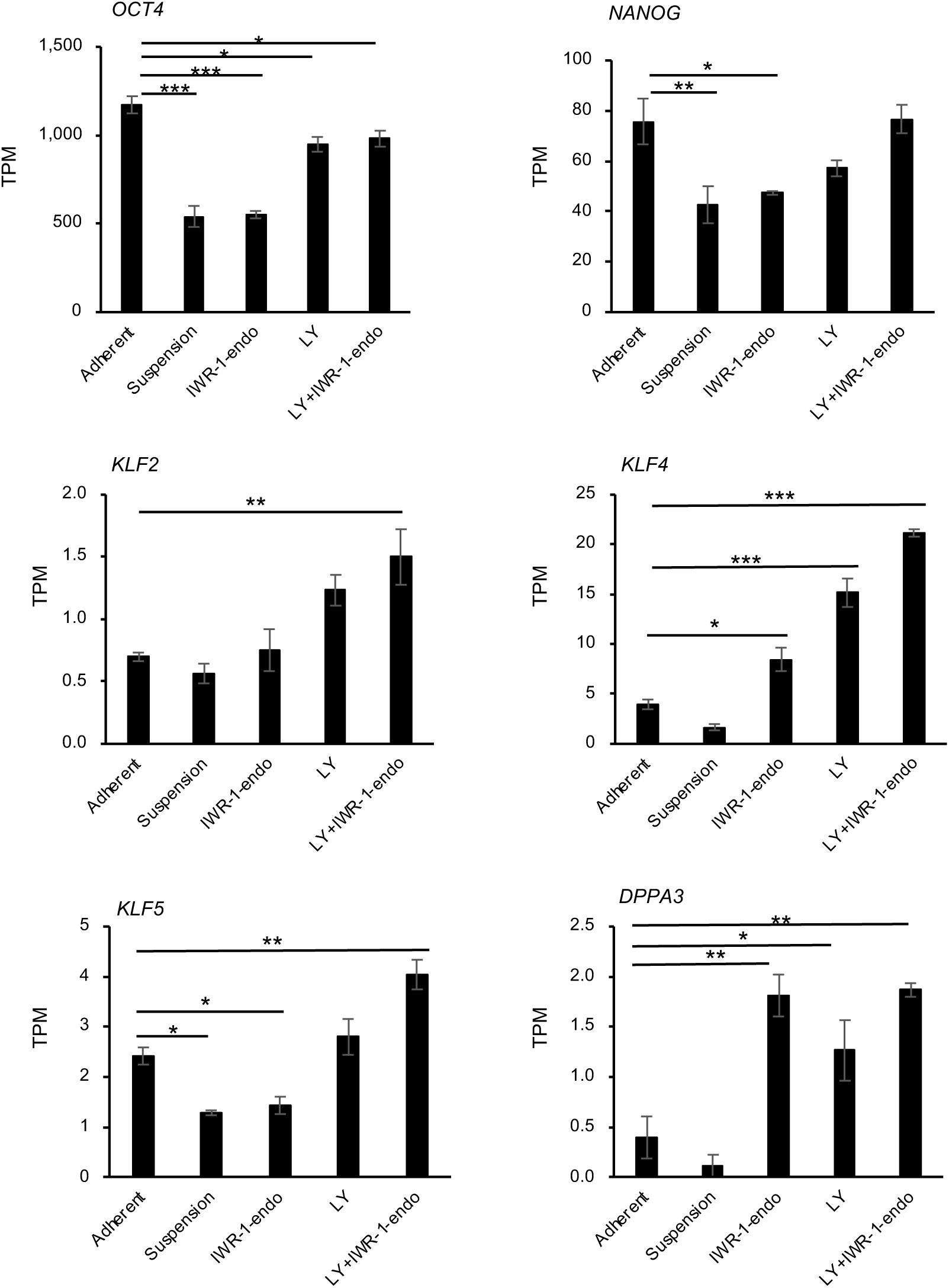
Comparison of expression on naïve markers between adherent and suspension-cultured hiPSCs. HiPSCs (WTC11 line) were culture in continuous suspension culture in StemFitAK02N in the presence of IWR-1-endo, LY333531, or both. RNA-seq was performed using day 10 samples. The gene expression of naïve pluripotency markers (*OCT4, NANOG, KLF2, KLF4, KLF5,* and *DPPA3*) was extracted based on the TPM (Transcripts per million) values. Bar graphs show the mean ± SE (n=3). P-values were statistically analyzed with Dunnett’s test to the samples of adherent conditions. P values < 0.05 were considered statistically significant. *, ** or *** in the graphs indicate P<0.05, P<0.01 or P<0.001, respectively.

**Figure 6—figure supplement 1.**
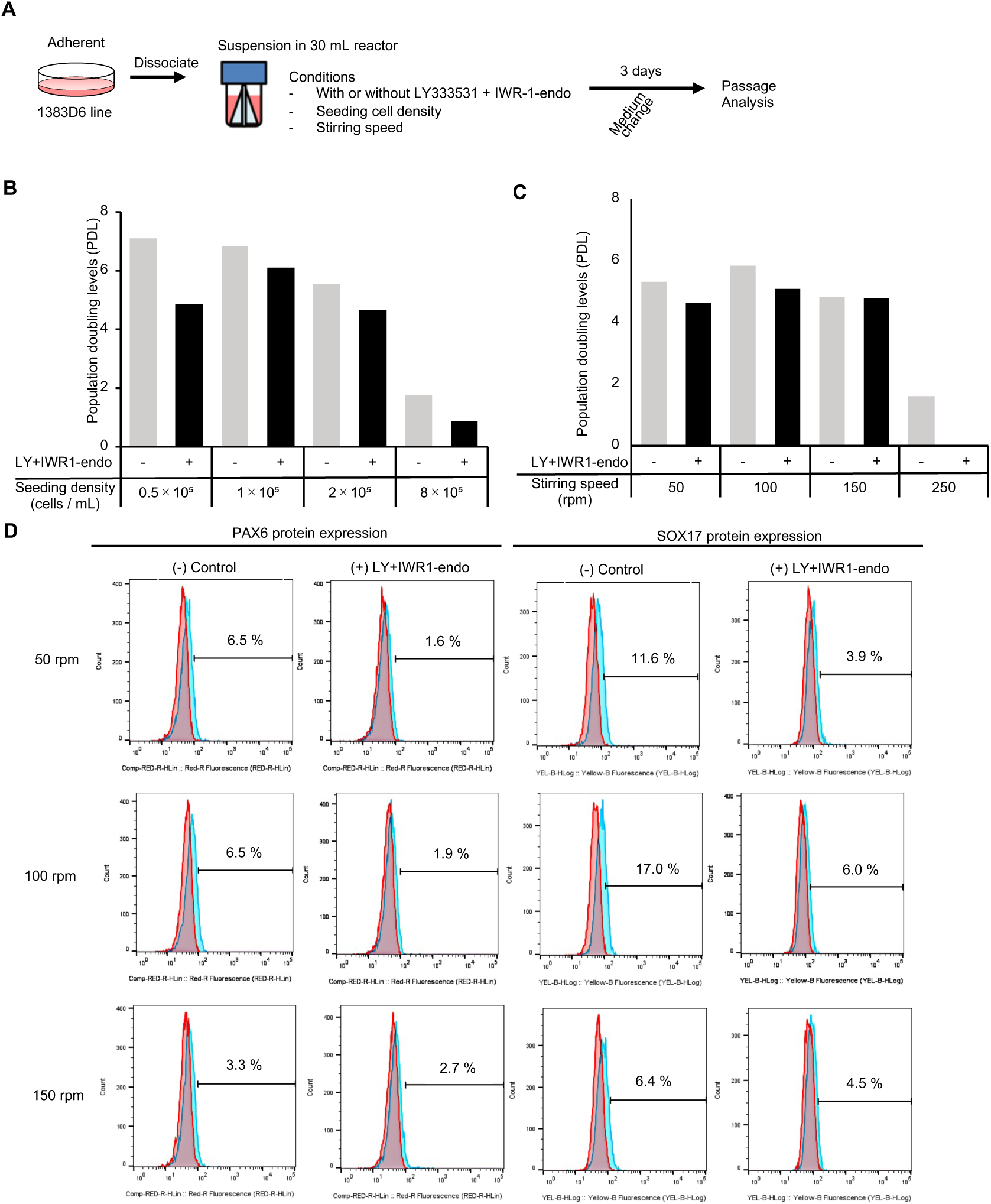
Suspension-culture of hiPSCs supplemented with IWR-1-endo and LY333531 using bioreactors with continuous agitation. (A) Schematic representing suspension culture of hiPSCs in a 30 mL bioreactor. Healthy donor-derived hiPSCs (1383D6 lines) were grown with continuous agitation in the presence or absence of IWR-1-endo and LY333531. Passages were performed every 3 -4 days. (B) Graphs showing population doubling levels from the seeding cell numbers on day 3. (C) Flow cytometric analysis of of differentiation marker proteins, PAX6 and SOX17. Positive cells (%) are shown.

**Figure 6—figure supplement 2.**
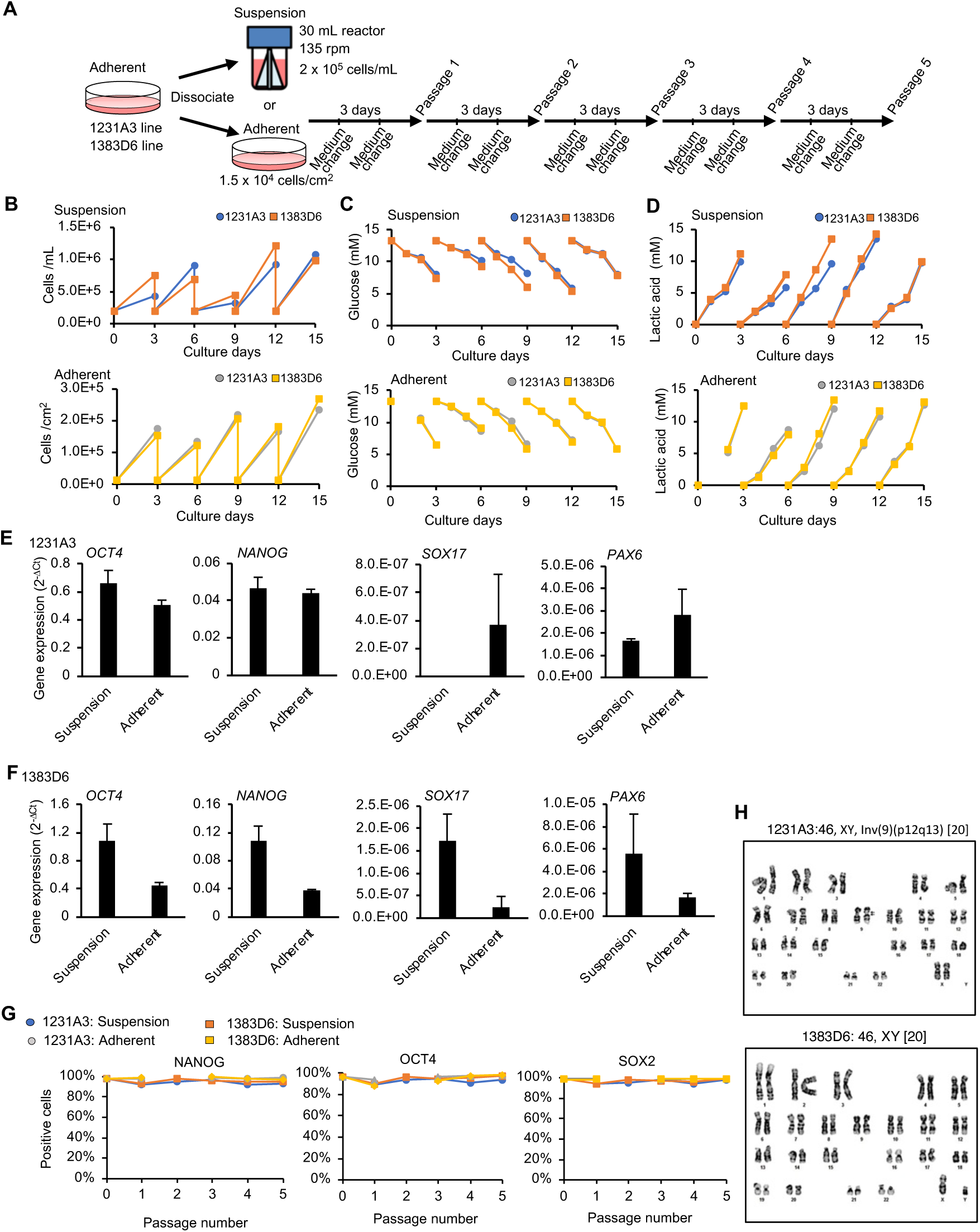
Suspension-culture of hiPSCs supplemented with IWR-1-endo and LY333531 using bioreactors with continuous agitation. (A) Schematic representing suspension culture of hiPSCs in a 30 mL bioreactor. Healthy donor-derived hiPSCs (1231A3 and 1383D6 lines) were grown with continuous agitation in the presence or absence of IWR-1-endo and LY333531. Passages were performed every three days. (B) Graphs showing cell numbers at each passage (n=1). Upper: suspension-culture of 1231A3 and 1383D6 lines. Lower: adherent-culture of 1231A3 and 1383D6 lines. (C) Graphs showing glucose concentration in the culture medium at each culture day. Upper: suspension-culture of 1231A3 and 1383D6 lines. Lower: adherent-culture of 1231A3 and 1383D6 lines. (D) Graphs showing lactic acid concentration in the culture medium at each culture day. Upper: suspension-culture of 1231A3 and 1383D6 lines. Lower: adherent-culture of 1231A3 and 1383D6 lines. (E and F) RT-qPCR analysis of hiPSCs cultured in 30 mL bioreactor or adherent conditions. cDNA samples were prepared from passage 3 and 5 of the 1231A3 in (E) and 1383D6 in (F). qPCR was performed for *OCT4* and *NANOG* (pluripotent), *CDX2* (mesoderm), *SOX17* (endoderm) and *PAX6* (neuroectoderm). The gene expression was normalized to *β-ACTIN*. Bar graph showing the average gene expressions of passage 3 and 5 (mean ± SE). (G) Flow cytometric analysis of suspension- or adherent-cultured 1231A3 and 1383D6 lines. Positive cells (%) of pluripotency marker proteins: NANOG, OCT4, and SOX2 are shown. (H) Karyotyping of hiPSCs cultured in 30 mL bioreactor. Cell samples were prepared on passage 5, and used for G-band karyotyping analysis. A short deletion on chromosome 9 of 1231A3 (indicated with arrow) was derived from the original line.

**Figure 8—figure supplement 1.**
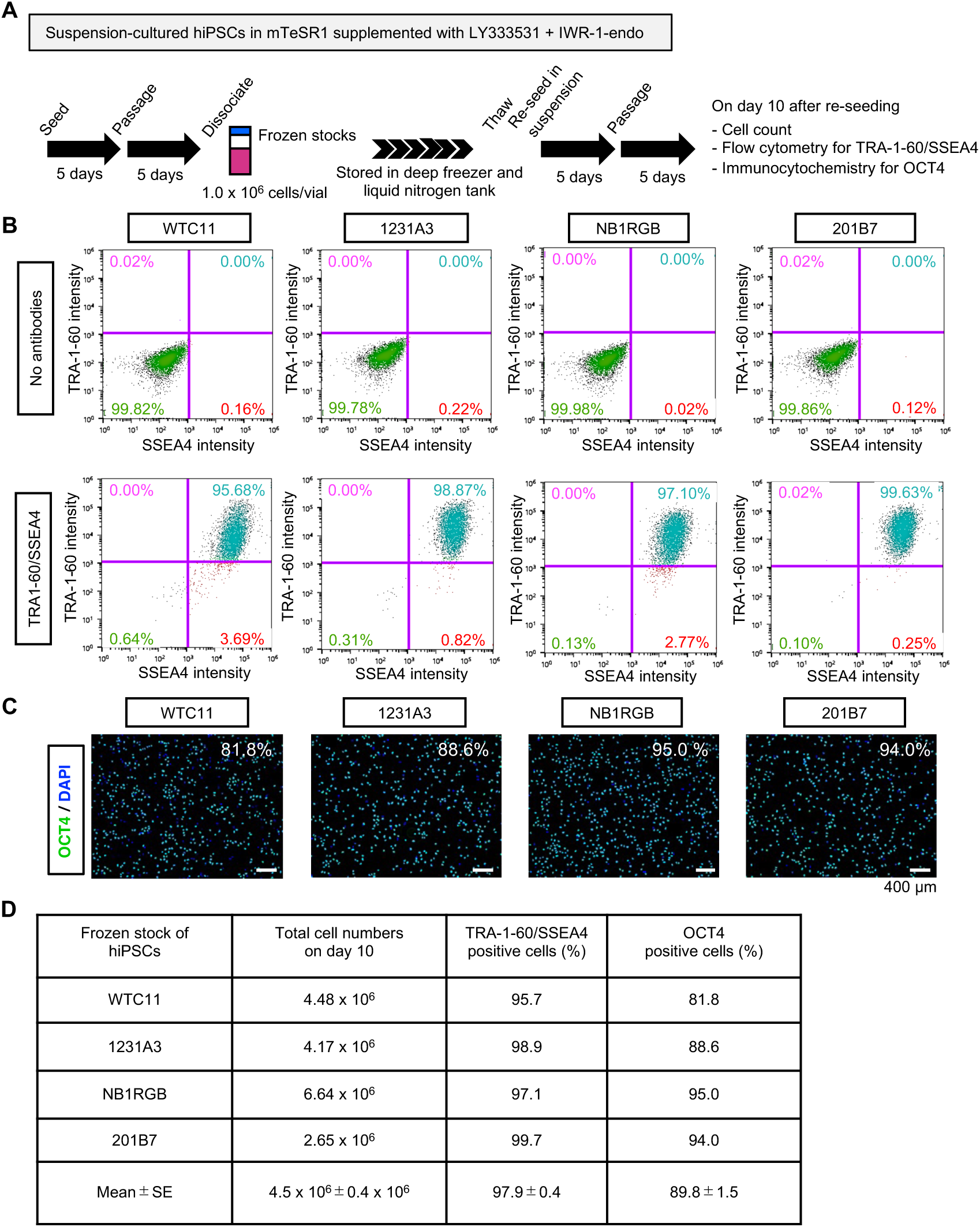
Re-suspension from frozen stocks of hiPSCs under suspension culture conditions supplemented with IWR-1-endo and LY333531 in mTeSR1 medium. (A) Schematics of re-suspension culture of frozen stocks of WTC11, 1231A3, HiPS-NB1RGB (NB1RGB) and 201B7 hiPSC lines. After suspension culture in mTeSR1 medium supplemented with IWR-1-endo and LY333531, expanded iPS cells were cryopreserved. Each frozen stocks were expanded again in the mTeSR1 medium supplemented with IWR-1-endo and LY333531 by repeating passage every 5 days. Characteristic analysis was performed on day10. (B) Representative flow cytometry of TRA-1-60 and SSEA4. (C) Representative immunocytochemistry of OCT4. DAPI indicates nuclear staining. Scale bars: 400 µm. (D) Summary of the characterization of re-suspension-cultured hiPSCs from frozen stocks. Total cell numbers on day 10, the ratio of double positive cells for TRA-1-60 and SSEA4 (%), and the ratio of OCT4-positive cells (%) are shown (mean ± SE; n=4).

**Figure 9—figure supplement 1.**
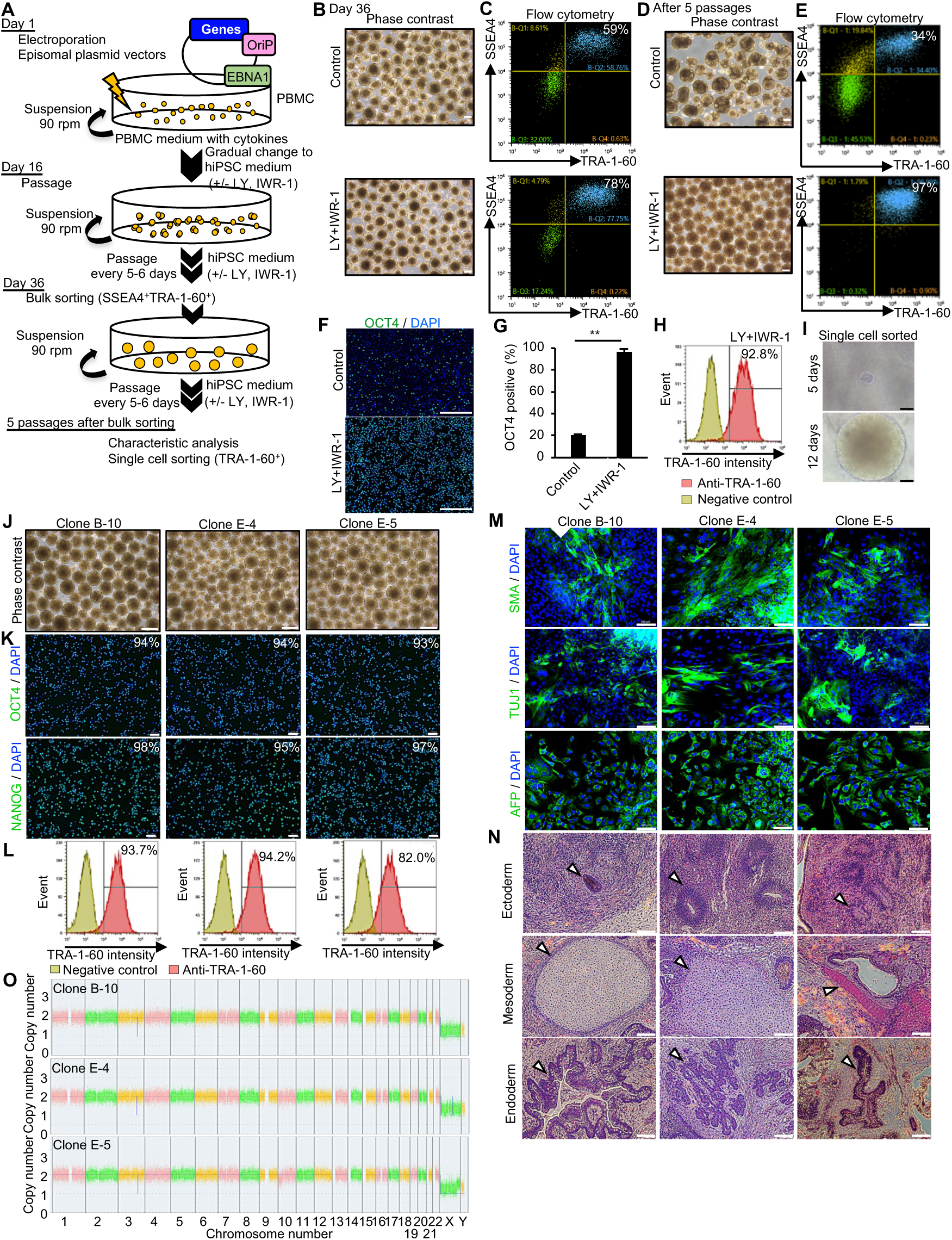
Establishment of hiPSCs in completed suspension conditions using episomal vectors. (A) Flow chart representing the establishment of hiPSCs from PBMCs by electroporation of episomal vectors. (B) Phase-contrast images of transfected cells on day 36 after electroporation of episomal vectors in the absence or presence of IWR-1-endo and LY333531. Scale bars: 400 µm. (C) Flow cytometry of TRA-1-60 and SSEA4. TRA-1-60 and SSEA4-psotive cells were sorted as the bulk fraction and further suspended in the absence or presence of IWR-1-endo and LY333531. The percentages of TRA-1-60 and SSEA4-psotive cells are shown in the right corner. (D) Phase-contrast images of suspension-cultured cell aggregates after bulk sorting on passage 5. Scale bars: 400 µm. (E) Flow cytometry of TRA-1-60 and SSEA4. The percentage of TRA-1-60- and SSEA4-psotive cells are shown in the right corner. (F) Immunocytochemistry of OCT4 on sorted-cells cultured in the absence (top) or presence (bottom) of IWR-1-endo and LY333531 for 5 passages. Scale bars: 400 µm. (G) Quantified data for OCT4-positive cell percentage. Bar graph indicating average OCT4-positive cell percentages (mean ± SE) (n=3). P-value was statistically analyzed by Student’s t-test. (H) Single-cell sorting and cloning of TRA-1-60 positive cells. Flow cytometry was performed for cell fraction that was cultured in the presence of IWR-1-endo and LY333531. TRA1-60-positive cells were collected as single cells in a 96-well plate and expanded in the culture medium supplemented with Y-27632, IWR-1-endo and LY333531. (I) Representative phase-contrast images of single-sorted cell aggregates on day 5 and 12. Scale bars, 400 µm. (J) Phase-contrast images of established clones (B-10, E-4, and E-5). Scale bars: 400 µm. (K) Immunocytochemistry of OCT4 and NANOG in these clones. The percentage of positive cell are shown. Scale bars: 100 µm. (L) Flow cytometry of TRA-1-60 in these clones. TRA-1-60 positive percentages are shown in the right corner. (M) Immunocytochemistry of differentiated cells in EBs from these clones. Anti-TUJ1, -SMA and -AFP antibodies were used. Scale bars: 100 µm. (N) Hematoxylin-Eosin staining of paraffin-embedded sections of teratoma derived from these clones. White arrowheads indicate representative tissue structures of the ectoderm, mesoderm, or endoderm. Scale bars, 100 µm. (O) Chromosomal copy numbers detected by CNV array analysis (Karyostat assay) of these clones.

**Supplementary Table 1.** The list of chemicals and cytokines used in the screening assay.

| Name | Effect | Manufacturer | Identifier | Concentration |
| --- | --- | --- | --- | --- |
| CHIR99021 | Wnt signal activator | Fujifilm Wako Pure Chemical | 034-23103 | 3 μM |
| IWP-2 | Wnt signaling inhibitor | Fujifilm Wako Pure Chemical | 034-24301 | 5 μM |
| IWR-1-endo | Wnt signaling inhibitor | Fujifilm Wako Pure Chemical | 037-25131 | 10 μM |
| Gö6976 | PKC ($\alpha$, $\beta$) inhibitor | Fujifilm Wako Pure Chemical | 071-06551 | 5 μM |
| Gö6983 | PKC ($\alpha$, $\beta$, $\gamma$, $\delta$) inhibitor | Fujifilm Wako Pure Chemical | 078-06443 | 5 μM |
| GF109203X | PKC ($\alpha$, $\beta$, $\gamma$) inhibitor | Tocris Bioscience | 0741 | 5 μM |
| Enzastaurin | PKC ($\alpha$, $\beta$, $\gamma$, $\epsilon$) inhibitor | Sigma-Aldrich | SML0762 | 5 μM |
| K-252a | PKC ($\mu$) inhibitor | Sigma-Aldrich | K1639 | 5 μM |
| LY333531<br>(Ruboxistaurin) | PKC ($\beta$) inhibitor | Sigma-Aldrich | SML0693 | 5 μM |
| Myricitrin | PKC ($\alpha$, $\epsilon$) inhibitor | Sigma-Aldrich | 91255 | 5 μM |
| Sotrastaurin | PKC ($\alpha$, $\beta$, $\delta$, $\epsilon$, $\eta$, $\theta$)<br>inhibitor | Cayman | 16726 | 5 μM |
| ZIP | PKC ($\zeta$) inhibitor | Tocris Bioscience | 2549 | 5 μM |
| Ro-32-0432 | PKC ($\alpha$, $\beta$, $\gamma$, $\epsilon$) inhibitor | Tocris Bioscience | 1587 | 5 μM |
| FK506 | Calcineurin inhibitor | Fujifilm Wako Pure Chemical | 063-06071 | 100 ng/mL |
| Cyclosporine A<br>(CsA) | Calcineurin inhibitor | Fujifilm Wako Pure Chemical | 031-24931 | 1 mg/mL |
| DAPT | γ-secretase inhibitor | Fujifilm Wako Pure Chemical | 043-33581 | 10 μM |
| BMP4 (Human,<br>recombinant) | BMP-SMAD signaling<br>activator | Fujifilm Wako Pure Chemical | 020-18851 | 40 ng/mL |
| LIF (Human,<br>recombinant) | LIF-STAT signaling<br>activator | Nacalai Tesque | NU0013-1 | 2000 units/mL |
| Valproic acid<br>(VPA) | Epigenetic modifier | Fujifilm Wako Pure Chemical | 227-01071 | 0.5 mM |
| Ascorbic acid | Epigenetic modifier | Nacalai Tesque | 13048-42 | 50 ng/mL |

**Supplementary Table 2.**
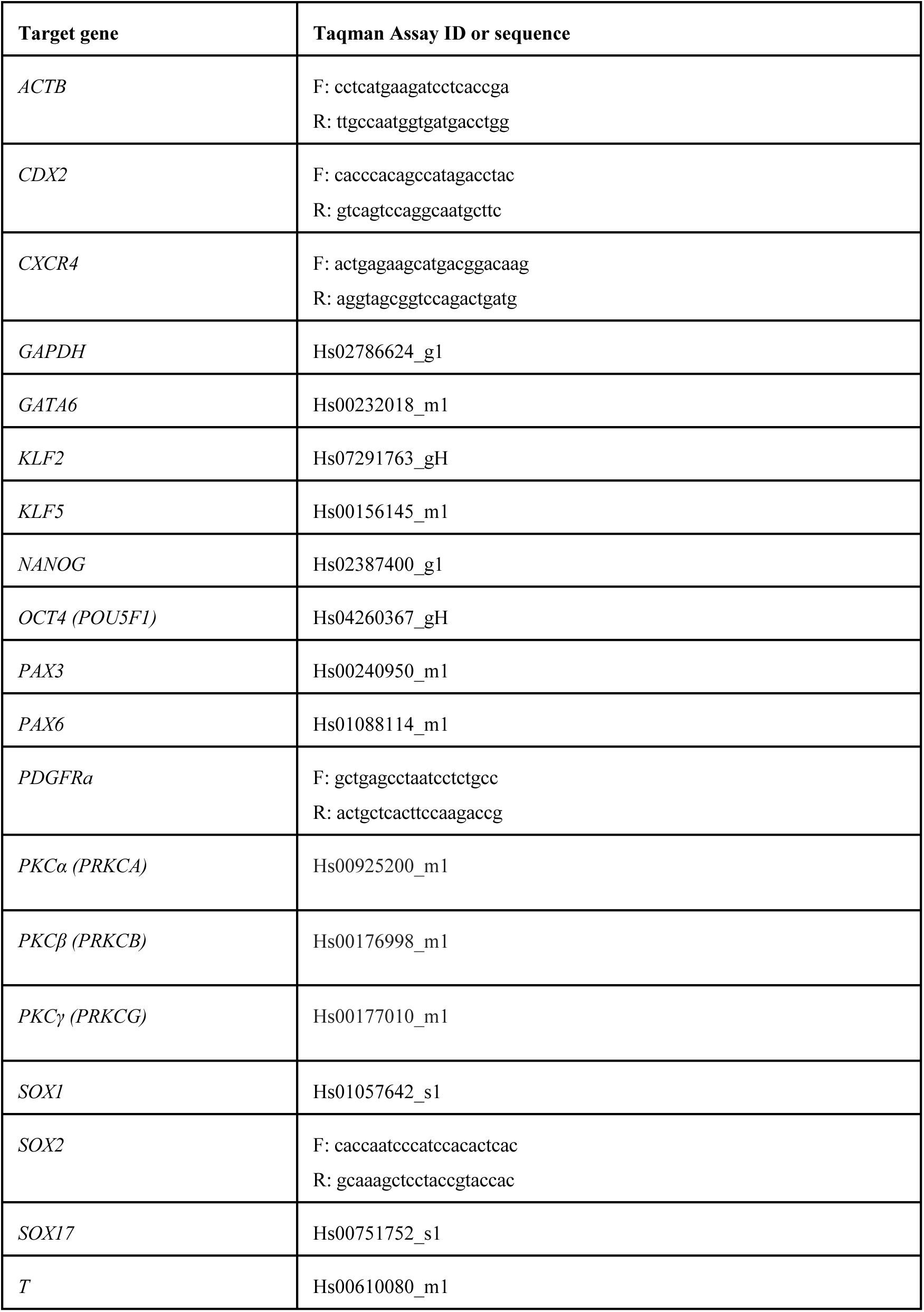
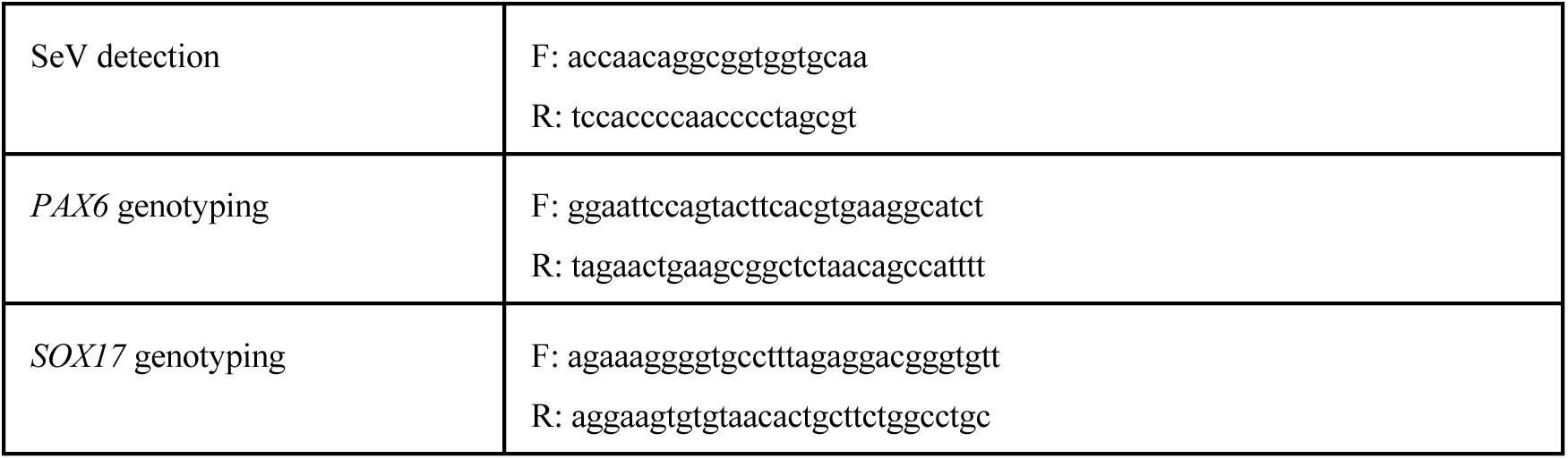
The list of Taqman probes and primer sets used in this study.

**Supplementary Table 3.** The list of primary antibodies used in this study.

| **Name** | **Host species** | **Dilutions or Concentrations** | **SOURCE** | **IDENTIFIER** |
| --- | --- | --- | --- | --- |
| Alpha-Fetoprotein (AFP) | Mouse | 1 µg/mL | R&D systems | Cat#MAB1368 |
| Alpha-SMA | Mouse | 1 µg/mL | R&D systems | Cat#MAB1420 |
| Cardiac Troponin T | Mouse | 10 µg/mL | Thermo Fisher Scientific | Cat#MA5-12960 |
| CORIN | Mouse | 10 µg/mL | Sigma-Aldrich | Cat#WH0010699M1 |
| GAPDH | Mouse | 1 µg/mL | R&D systems | Cat#MAB5718 |
| NANOG | Rabbit | 0.5 µg/mL | Reprocell | Cat#RCAB004P-F |
| OCT-3/4 | Mouse | 0.4 µg/mL | Santa Cruz | Cat#sc-5279 |
| Oct-4A (C30A3) Rabbit mAb (Alexa Fluor 488 Conjugate) | Rabbit | 1:30 | Cell Signaling Technology | Cat# 5177S |
| PAX6 | Rabbit | 1:400 (Immunocytochemistry) | MBL | Cat#PD022 |
| PAX6 | Sheep | 5 µg/mL (Wes) | R&D Systems | Cat#AF8150 |
| Anti-PKC beta1 + PKC beta2 (phospho T641) | Rabbit | 1:50 | Abcam | Cat#ab194749 |
| SOX17 | Goat | 0.4 µg/mL (Immunocytochemistry) | R&D systems | Cat#AF1924 |
|  |  | 1 µg/mL (Wes) |  |  |
| FITC Mouse anti-SSEA-4 | Mouse | 1:5 | BD Bioscience | Cat#560126 |
| APC anti-human SSEA4 | Mouse | 1:20 | BioLegend | Cat#330418 |
| Alexa Fluor 488 anti-human TRA1-60-R | Mouse | 1:20 | BioLegend | Cat#330614 |
| Alexa Fluor 488 Mouse anti-Human TRA-1-60 Antigen | Mouse | 1:10 | BD Bioscience | Cat#560173 |
| TUJ1 | Mouse | 1 µg/mL | R&D systems | Cat#MAB1195 |

**Supplementary Table 4.** The list of secondary antibodies used in this study.

| Name | Host species | Dilutions | SOURCE | IDENTIFIER |
| --- | --- | --- | --- | --- |
| Anti-Mouse IgG (H+L) Highly Cross-Adsorbed Secondary Antibody, Alexa Fluor Plus 488 | Donkey | 1:1000 | Thermo Fisher Scientific | Cat#A-21202 |
| DyLight 488 anti-rabbit IgG (minimal x-reactivity) | Donkey | 1:1000 | BioLegend | Cat#406404 |
| Anti-Mouse IgG (H+L) Highly Cross-Adsorbed Secondary Antibody, Alexa Fluor Plus 555 | Donkey | 1:1000 | Thermo Fisher Scientific | Cat#A-31570 |
| Anti-Rabbit IgG (H+L) Cross-Adsorbed Secondary Antibody, Alexa Fluor 555 | Goat | 1:1000 | Thermo Fisher Scientific | Cat#A-21428 |
| Anti-Mouse IgG (H+L) Cross-Adsorbed Secondary Antibody, Alexa Fluor 488 | Goat | 1:1000 | Thermo Fisher Scientific | Cat#A-11001 |
| Donkey Anti-Sheep IgG HRP Affinity Purified PAb | Sheep | 1:50 | R&D systems | Cat# HAF016 |

**Supplementary Table 5.** Software used in this study.

| Name | Source | URL |
| --- | --- | --- |
| clusterProfiler (v3.16.1) | R package | https://bioconductor.org/packages/release/bioc/html/clusterProfiler.html |
| edgeR (v3.30.3) | R package | https://bioconductor.org/packages/release/bioc/html/edgeR.html |
| R (version4.1.0) | R package | https://www.r-project.org/ |
| CLC Genomics Workbench (v20) | QIAGEN | https://digitalinsights.qiagen.com/products-overview/discovery-insights-portfolio/analysis-and-visualization/qiagen-clc-genomics-workbench/ |
| SH800S cell sorter and flow cytometry software | Sony | https://www.sonybiotechnology.com/us/instruments/sh800s-cell-sorter/software/ |
| Compass software for Simple Western | Protein Simple | https://www.biotechne.com/resources/instrument-software-download-center/compass-software-simple-western |
| BZ-X800 microscope analysis software | Keyence | https://www.keyence.com/landing/microscope/lp_fluorescence.jsp |
| QuantStudio 3 and 5 Real-Time PCR System Software | Thermo Fisher Scientific | https://www.thermofisher.com/jp/ja/home/global/forms/life-science/quantstudio-3-5-software.html |
| Chromosome Analysis Suite (ChAS) and Affymetrix GeneChip Command Console software programs | Thermo Fisher Scientific | https://www.thermofisher.com/jp/ja/home/life-science/microarray-analysis/microarray-analysis-instruments-software-services/microarray-analysis-software/chromosome-analysis-suite.html |
| CellSens microscope software | Evident (Olympus) | https://www.olympus-lifescience.com/en/support/downloads/ |

## Notes

### Summary of Updates

We have completed our revision to meet all the helpful comments by the reviewers. Major points are below. 1. We tested several cell seeding densities and several stirring speeds with or without WNT/PKCβ inhibitors (Figure 6-figure supplement 1). 2. We have performed additional experiments using conventional media, mTeSR1 (Stem Cell Technologies, Vancouver, Canada), comparing with the adherent feeder-free culture system in four different hiPSC lines simultaneously (Figure 4 - figure supplement 2). 3. We have performed additional experiments on hiPSC maintenance across 5 hiPSC lines in suspension culture using StemFit AK02N medium simultaneously (Figure 3C - E). 4. We found that the expression of naive pluripotency markers, KLF2, KLF4, KLF5, and DPPA3, were up-regulated in the suspension conditions treated with LY333531 and IWR-1-endo while the expression of OCT4 and NANOG was at the same levels (Figure 5-figure supplement 2). 5. We added the data on the cloning efficiency, which are compared with adherent cultures (Figure 7B). 6. We have added more detailed explanation in the Discussion section

